# The microtubule-associated protein SlMAP70 interacts with SlIQD21 and regulates fruit shape formation in tomato

**DOI:** 10.1101/2022.08.08.503161

**Authors:** Zhiru Bao, Ye Guo, Yaling Deng, Jingze Zang, Junhong Zhang, Bo Ouyang, Xiaolu Qu, Katharina Bürstenbinder, Pengwei Wang

## Abstract

The shape of tomato fruits is closely correlated to microtubule organization and the activity of microtubule associated proteins (MAP), but insights into the mechanism from a cell biology perspective are still largely elusive. Analysis of tissue expression profiles of different microtubule regulators revealed that functionally distinct classes of *MAPs* are highly expressed during fruit development. Among these, several members of the plant-specific *MAP70* family are preferably expressed at the initiation stage of fruit development. Transgenic tomato lines overexpressing SlMAP70 produced elongated fruits that show reduced cell circularity and microtubule anisotropy, while SlMAP70 loss-of-function mutant showed an opposite effect with flatter fruits. Microtubule anisotropy of fruit endodermis cells exhibited dramatic rearrangement during tomato fruit development, and SlMAP70-1 is likely implicated in cortical microtubule organization and fruit elongation throughout this stage by interacting with SUN10/SlIQD21a. The expression of *SlMAP70* (or co-expression of *SlMAP70* and *SUN10/SlIQD21a*) induces microtubule stabilization and prevents its dynamic rearrangement, both activities are essential for fruit shape establishment after anthesis. Together, our results identify SlMAP70 as a novel regulator of fruit elongation, and demonstrate that manipulating microtubule stability and organization at the early fruit developmental stage has a strong impact on fruit shape.

## Introduction

Tomato (*Solanum lycopersicum*) fruits are composed of pericarp tissues with at least two carpels, and placental tissues surrounding the seeds. The development from flower bud primordia to mature fruits involves four important stages: flower initiation and development, intensive cell division, rapid cell expansion, and fruit ripening (Tanksley, 2004). Changes in cell division and/or expansion during early fruit development stages are believed to have direct impacts on fruit yield and quality. In tomato, the genetic basis of fruit shape formation has been assigned to several genes that group into two categories: genes that regulate the number of fruit locules such as *FASCIATED* (*FAS*) and *LOCULE NUMBER* (*LC*) (Chu et al., 2019; Muños et al., 2011), and genes which regulate fruit cell division or elongation such as *SUN* and *OVATE* (Liu et al., 2002; Xiao et al., 2008).

Recent findings suggest that microtubules are important regulators of fruit morphogenesis (Bao et al., 2021; Lazzaro et al., 2018). Several key protein families, such as SUN, OFP (OVATE Family Protein), TRM (TONNEAU1 Recruiting Motif) and KTN1 (katanin), have been implicated in the regulation of fruit shape/ and size in plants (Guo et al., 2020; Snouffer et al., 2020; Wang et al., 2021). In tomato, OFPs regulate cell division patterns during fruit development by interacting with TRM proteins (Wu et al., 2018), which are essential for the recruitment of TTP (TON1– TRM–PP2A) complex to microtubules and the assembly of the preprophase band (PPB) during cell division (Schaefer et al., 2017; Spinner et al., 2013).

Tomato SUNs, also known as IQ67-domain (IQD) proteins, comprise a family of microtubule associated proteins (MAPs). SUN1/SlIQD12a is encoded by a quantitative locus, which was first identified in the tomato *sun* mutant. The sun locus contains a retrotransposon-mediated gene duplication resulting in increased *SUN1* expression and elongated fruits (Huang et al., 2013; Xiao *et al*., 2008). The cellular functions of SUN likely are conserved in different plant species. Mis-expression of several *AtIQDs* in *Arabidopsis thaliana* correlates with altered growth and re-organized cortical microtubules arrays (Bürstenbinder et al., 2017; Kumari et al., 2021; Mitra et al., 2019; Zang et al., 2021). Such phenomenon also found in the grain/fruit organ morphogenesis of rice (Duan et al., 2017; Liu et al., 2017; Yang et al., 2020), cucumber and watermelon species (Dou et al., 2018; Pan et al., 2017; Pan et al., 2020). Collectively, these findings imply that modulating *SUN* expression and activity changes organ shapes, and that different SUN/IQD members may have functional redundancy in this process.

At the molecular level, SUN/IQDs are proposed to function as scaffolding proteins in cell signaling (Abel et al., 2013; Bürstenbinder et al., 2013). *Arabidopsis* IQDs have been reported to interact with various MAPs and MAP-related proteins, including the microtubule-severing enzyme KTN1 (Li et al., 2021), Kinesin Light Chain-Related proteins (KLCRs) (Bürstenbinder *et al*., 2013; Liu et al., 2016; Yang et al., 2022; Zang *et al*., 2021), Phragmoplast Orienting Kinesins (POKs) and Pleckstrin-Homology GTPase-Activating proteins (PHGAPs) (Kumari *et al*., 2021) to regulate apical hook formation, cellulose deposition, cell morphogenesis, cell division-plane orientation and establishment, respectively. These observations support the idea that distinct classes of MAPs regulate cell anisotropic growth, division patterns, and overall organ shape via differential microtubule modulating activities. These activities likely are fine-tuned by differential and dynamic interactions between functionally distinct MAPs and MAP-related proteins. However, which MAPs are essential during tomato fruit morphogenesis, and whether similar mechanisms are found in fruit morphogenesis remain to be confirmed.

MAP70 proteins (MAP70s) are a plant-specific class of MAPs (Korolev et al., 2007; Korolev et al., 2005; Pesquet et al., 2011) that are divided into two major clades, the MAP70-1 and -5 clades (Pesquet *et al*., 2011). In *Arabidopsis*, MAP70-5 regulates cortical microtubule organization within the inner side of endodermal cells via its bundling effect, changing the endodermal cell wall to facilitate lateral root morphogenesis (Stöckle et al., 2022) However, MAP70-1 and MAP70-5 isoforms only share a maximum similarity of 37% in protein sequence (Pesquet *et al*., 2011), suggesting that MAP70-1 may have different functions from MAP70-5.

Here, we show that *MAP70-1* from tomato is highly expressed at the early fruit developmental stages, and identify MAP70 family members as novel regulators for cortical microtubule arrays patterning and fruit shape formation in tomato. Mutants with knock-down (or knock-out) expression of *SlMAP70-1*, *SlMAP70-2* and *SlMAP70-3* display round cotyledons and flat fruits, while overexpressing *SlMAP70-1* induces the opposite effect. The altered fruit shape is mainly due to changes in cell shape and cortical microtubule arrays starting from ovary development. A yeast-two-hybrid test identified SUN10/SlIQD21a as a novel MAP70 interactor. In the presence of SlMAP70 and SUN10/SlIQD21a, the average microtubule length increases significantly, and tend to form whorl-like arrays in *N. benthamiana* leaf epidermal cells. Overexpression of *SUN10/SlIQD21a* induces elongated fruits, similar to *SlMAP70-1* over-expression lines, which is further enhanced upon overexpression of *SUN10/SlIQD21a* and *SlMAP70-1* together. These results identify MAP70s as novel regulators of fruit shape that function together with IQDs, and provide novel mechanistic insights and in-depth subcellular observations of microtubule organization underlying fruit shape regulation in tomato.

## Results and Discussions

### *SlMAP70-1* is highly expressed at the early stage of tomato fruit development

Microtubule activity is essential for cell morphogenesis and organ shape formation (Buschmann and Müller, 2019; Eng and Sampathkumar, 2018). To identify novel microtubule-related candidates involved in tomato fruit size regulation or shape formation, we analyzed the gene expression of multiple microtubule-associated proteins (MAPs) in tomato reproductive organs during fruit development. We found that several MAP genes from the Spiral1 (SPR1), SPR2, TONNEAU2 (TON2), Wave-Dampened2 (WVD2) and MAP70 families are highly expressed (Figure S1), suggesting these genes may be involved in fruit development. MAP70 was chosen for further characterization, as its homologues also exhibit very high-level of expression in fruits of other species (e.g. cucumber) (Jiang et al., 2015). BLAST search identified five MAP70 homologues in the tomato genome (version SL4.0, https://solgenomics.net/). Phylogenetic and expression analysis indicated that SlMAP70-1, -2, -3 belong to the MAP70-1 clade, which are highly expressed in fruits; whereas SlMAP70-4 and -5 belong to the MAP70-5 clade that is preferably expressed in roots (Figure 1 a-b).

To investigate the function of the tomato MAP70 family, we cloned all five tomato MAP70 members, generated GFP fusion constructs, and analyzed their subcellular localization in transient expression assays in *N. benthamiana*. In interphase leaf epidermal cells, all GFP-SlMAP70 proteins localized to the microtubules, as evidenced by co-alignment with the microtubule marker RFP-Tubulin5α (RFP-TUA5; Figure S2). To test if SlMAP70s are also present at mitotic microtubule arrays, we induced cell division by co-expression with cyclinD3;1 (Xu et al., 2019). During cell division, most GFP-SlMAP70 co-localized with microtubules at the preprophase band, spindle and phragmoplast, except for GFP-SlMAP70-4 which also labels the cell plate in telophase and the nucleus in preprophase (Figure S2). These data establish tomato SlMAP70s as *bona fide* MAPs that localize to diverse microtubule arrays during interphase and cell division. As most SlMAP70 members share high sequence similarity, SlMAP70-1, which has the strongest expression at the fruit initiation stage (Figure 1b), was selected for further functional studies.

**Figure 1.**
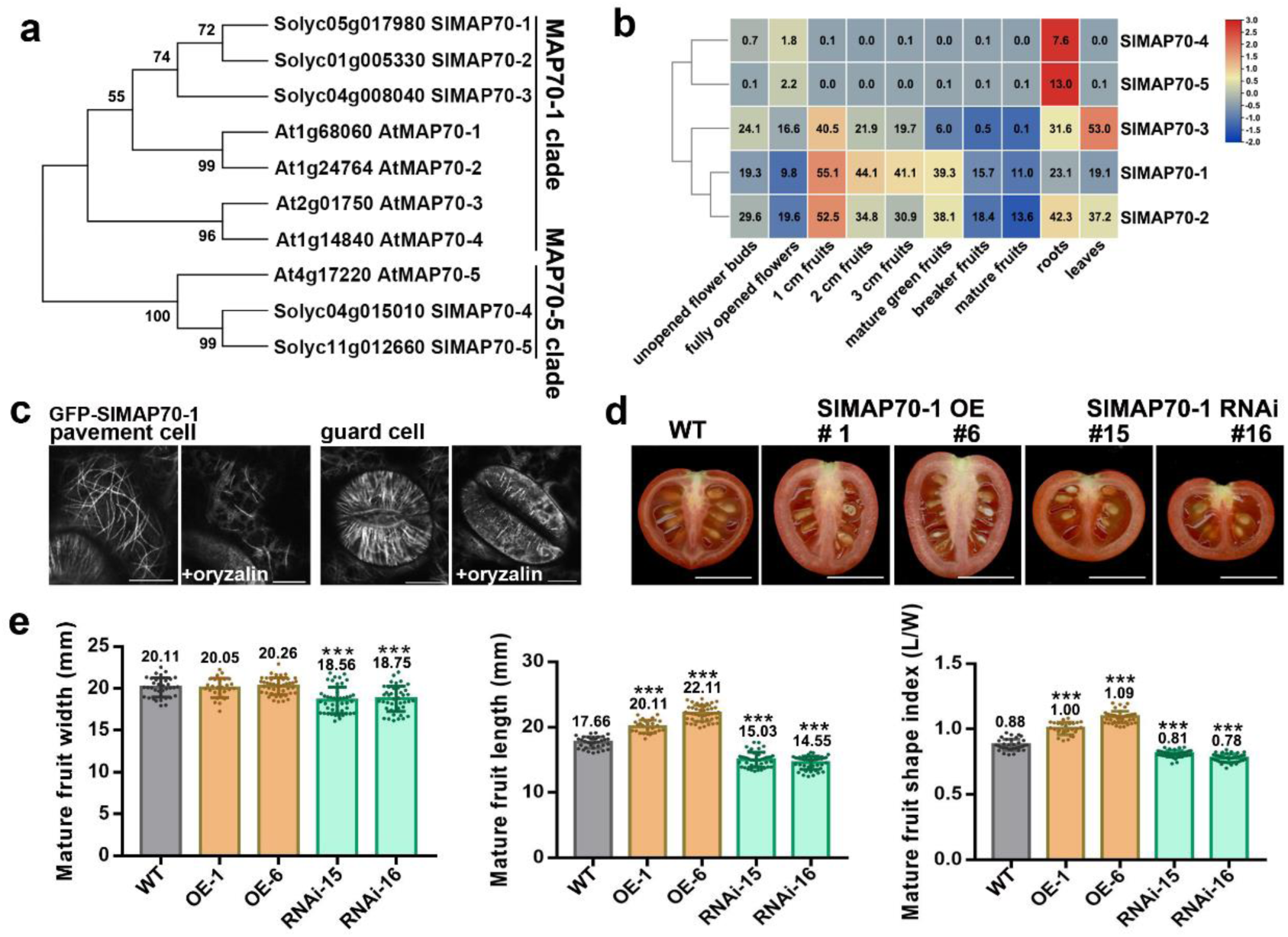
The SlMAP70 proteins localized to the microtubules and affect fruit morphogenesis. **(a)** Phylogenetic tree of *MAP70* gene family in tomato and *Arabidopsis*, numbers at the tree forks indicated bootstrap values. **(b)** Hierarchical clustering of expression values of *SlMAP70* genes in different tissues. Red indicates high expression level; blue, low expression level. Genes of MAP70-1 clade expressed higher in early fruit stage (1cm-size fruit), and MAP70-5 clade mainly expressed in root tissue. **(c)** Subcellular localization of GFP-SlMAP70-1 in tomato leaf pavement cells. Oryzalin treatment assays indicated that SlMAP70-1 labeled filaments are microtubules. Bars: 10 μm. **(d-e)** Effects of *SlMAP70-1* expression on fruit shape. Bars: 1 cm. Overexpression of *SlMAP70-1* under the control of the 35S promoter (*SlMAP70-1*-OE lines) enhances fruit longitudinal growth, while down-expression by RNA interference (*SlMAP70*-1-RNAi lines) showed flatter fruit shape. The error bars and asterisks in the graph indicate the standard errors and significant differences compared with wilt-type evaluated by Tukey’s test (p < 0.001). n > 25.

### Tomato fruit elongation and cell morphogenesis are controlled by the activity of SlMAP70

To functionally characterize SlMAP70-1, a wild type tomato (MicroTom) plants were transformed using either a *Pro35S:GFP-SlMAP70-1* construct or an RNA interference construct that was targeted to the *SlMAP70-1* gene (Figure S3a). In transgenic tomato *Pro35S:GFP-SlMAP70-1* lines, GFP-SlMAP70-1 localized to filaments that were sensitive to oryzalin treatment, a drug preventing microtubule polymerization (Morejohn et al., 1987), confirming microtubule localization (Figure 1c). Plants with elevated levels of *SlMAP70-1* exhibited elongated fruit morphology, while fruits with knocked-down expression of *SlMAP70-1* were flatter (Figure 1d). The phenotype was consistently observed in multiple independent transgenic lines (Figure 1d-e and S3b-c). Quantitative and statistical analysis revealed that fruit length was increased in *SlMAP70-1* over-expressing lines (denoted as OE), and oppositely, in *SlMAP70-1* RNAi lines fruit length and fruit width were reduced (Figure 1 e), resulting in increased and reduced fruit shape indices (as indicated by the length/width ratio), respectively. *SlMAP70* RNAi construct was also transformed into another tomato cultivar, AC (Ailsa Craig). As observed in MicroTom, transgenic AC fruits were flatter than the wild type fruits (Figure S4 a-b). These data, indicate that the function of SlMAP70 during fruit shape formation is likely conserved in different tomato varieties.

As a high sequence similarity is found among the MAP70-1 clade, the expression of *SlMAP70-2*, *3* was also slightly reduced in *SlMAP70-1* RNAi lines (Figure S3 a-b), suggesting that the RNAi construct may interfere other *SlMAP70* members that are likely functionally redundant in fruit development. To address potential functional redundancies, loss-of-function mutants of different *SlMAP70* members were generated using the CRISPR/Cas9 technique, and different combinations of *slmap70 crispr (cr)* mutants were obtained (Figure S4c, e-h). Loss of *SlMAP70-1* and *SlMAP70-2* in the *slmap70-1 cr* and *slmap70-2 cr* single mutants, respectively, resulted in reduced shape indexes, and the reduction in fruit length was further enhanced in *slmap70-1/2 cr* double mutants. On the other hand, *slmap70-4/5 cr* double mutants did not exhibit significant changes in fruit phenotypes (Figure S4 c-d), indicating that SlMAP70-1 and -2 are likely the dominant players in regulating tomato fruit shape and elongation.

In addition to changes in fruit shape, cell morphological changes were observed in other tissues of *SlMAP70-1* transgenic plants (Figure S5a). Hypocotyls were elongated in *SlMAP70-1* OE lines but shortened in *SlMAP70-1* RNAi lines (Figure S5b), and cell shape complexity (indicated by cell circularity) was increased and reduced in leaf epidermis pavement cells and mature fruit endocarp cells of *SlMAP70-1* OE lines and *SlMAP70-1* RNAi lines, respectively (Figure 2a, b). These results suggested that the function of SlMAP70 in cell and tissue morphogenesis is likely conserved in different cell types.

**Figure 2.**
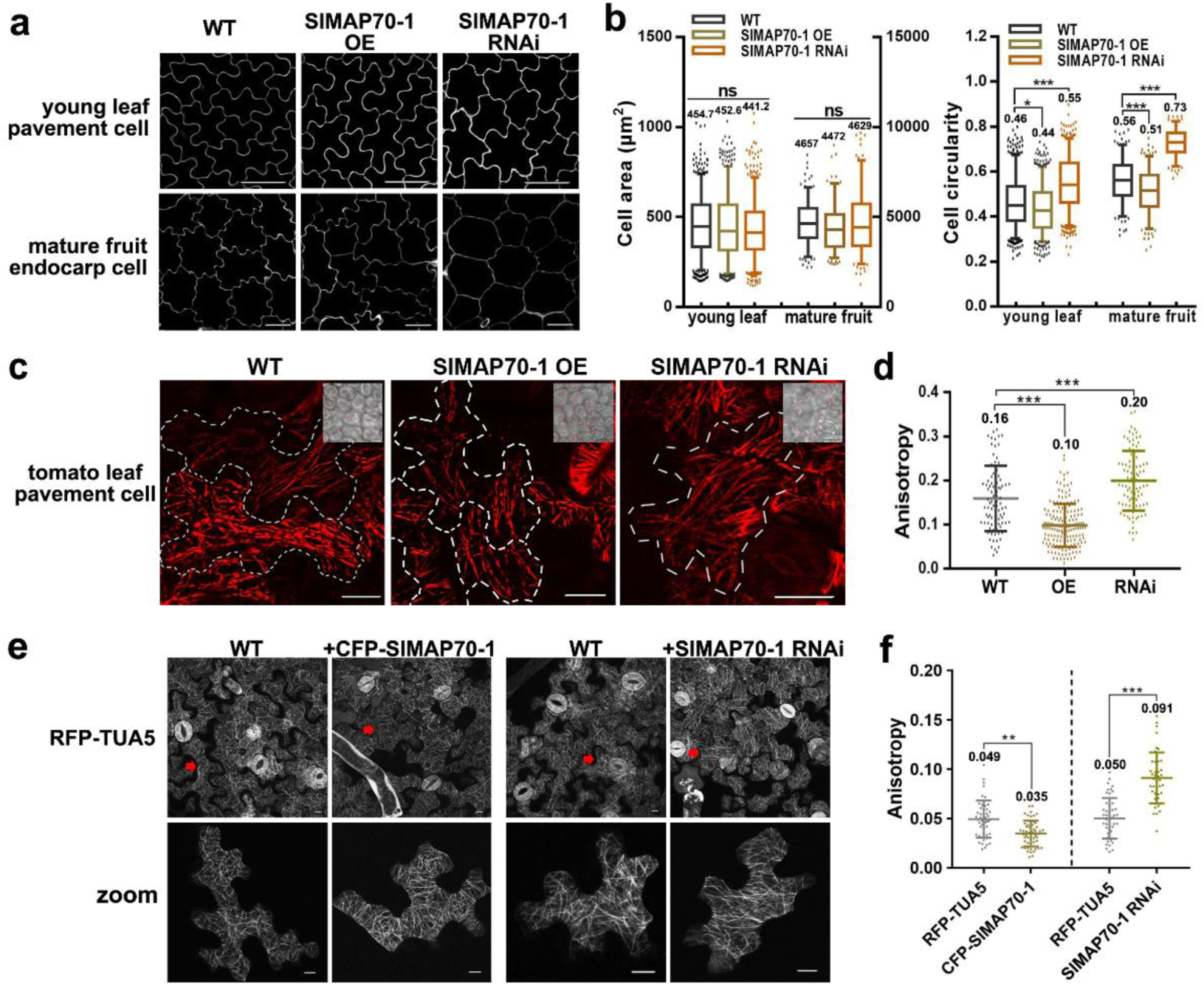
Analysis of microtubule arrangement and cell shape in *SlMAP70* misexpression plants. **(a)** Leaf pavement cells of 15-day-old seedlings and endodermis cells of mature fruits. Cell contours were visualized by PI staining. Bars: 50 μm for endocarp PCs, Bars: 25 μm for young leaf PCs. **(b)** Quantification of cell size (area) and shape complexity (circularity). Overexpression of *SlMAP70-1* caused more irregular cell morphology, and down-expression of *SlMAP70* lead to more circular cells. Similar results were observed in leaf pavement and fruit endocarp tissue. Cells from five confocal images of three different plants were analyzed (n > 100). ns: no significance; *, p<0.05; **, p<0.01; ***, p<0.001 by Tukey’s test. **(c)** Immunofluorescence of tomato young leaf epidermal cells with a tubulin antibody. Dotted lines mark the cell outline. Bars: 10 μm. **(d)** Quantitative analysis of microtubule array anisotropy in young leaf of different SlMAP70 genotypes. Anisotropy was measured by FibrilTool. Center line and error bars indicate means ± SD (n > 50 cells). ***, p < 0.001 by Tukey’s test. **(e)** Cortical MTs of *N. Benthemiana* leaf pavement cells with overexpressed CFP-SlMAP70-1, or with knock-down expression of *NbMAP70*. bars: 10 μm. **(f)** Quantitative analysis of MT array anisotropy in (e) and (f) by the ImageJ FibrilTool plugin. Center line and error bars indicate means ± SD (n = 50). **, p < 0.01 and ***, p < 0.001 by Tukey’s test.

### SlMAP70 proteins regulate microtubule patterning

The arrangement of microtubules has a direct impact on cell expansion and anisotropy of growth. We thus suspected that the altered cell size, cell circularity (Figure 2 a-b) and organ shape (Figure 1d-e) are likely to link to changes in microtubule organization regulated by SlMAP70. To test this hypothesis, we immuno-labeled the cortical microtubules of leaf epidermal cells from the wild-type tomato, *SlMAP70-1* OE and the RNAi lines (Figure 2c). To assess whether microtubules are ordered, anisotropy scores were calculated. The score of 0 represents no order (all filaments arranged at different directions) and 1 represents perfect order (all filaments at the same direction)(Boudaoud et al., 2014). The average anisotropy score in wild type plant was significantly higher than that in *SlMAP70-1* OE lines, but lower than that in *SlMAP70-1* RNAi lines (Figure 2d), suggesting a function of SlMAP70 in microtubule arrangement. Unfortunately, similar subcellular studies could not be conducted in fruits, as it is difficult to fix cells from fruits, and preserving the subcellular structure of cells with high-water content is extremely difficult. However, we speculated that similar changes in microtubule organization are likely occurring in fruits, as the endocarp cell morphology also exhibit similar alterations as observed in leaf and cotyledon cells (Figure 2 a-b, Figure S5).

In addition, similar results were observed in *N. benthamiana* pavement cells stably transformed with a microtubule marker, RFP-TUA5 (Figure 2 e-f). Disordered microtubules were found in *SlMAP70-1* OE cells, whereas an increase in ordered microtubule arrangements was found in cells with reduced *MAP70* level (upon over-expressing of the *SlMAP70-1* RNAi construct that likely cross-reacts with *MAP70* genes in *N. benthamiana*). The over-expression of CFP-SlMAP70-1 and knock-down effects of NbMAP70 were confirmed by western blot using a SlMAP70-1 antibody that recognized its *N. benthamiana* homologue (Figure S6 a-c). Taken together, these results indicate that SlMAP70 is required to regulate microtubule organization, an important feature for cell shape establishment.

### SlMAP70s interact with SlIQD21 proteins

To characterize the role of SlMAP70 proteins in fruit morphology, a yeast two-hybrid screen was performed using SlMAP70-1 as bait. As potential interaction partner, SlIQD21a (also named SUN10 from the tomato genome annotation) was identified. Public expression data (The tomato genome sequence provides insights into fleshy fruit evolution, 2012) indicate that *SUN10/SlIQD21a* is expressed at a high levels in early developing fruits and young leaves, similar to the expression pattern of *SlMAP70-1* (Figure 1b, Figure S7a). Analysis of SUN10/SlIQD21a subcellular localization showed that GFP-SUN10/SlIQD21a localizes to cortical microtubules, as indicated by co-localization with RFP-TUA5 (Figure S7b). Co-expression of RFP-SUN10/SlIQD21a and GFP-SlMAP70-1 in *N. benthamiana* leaf pavement cells showed clear co-localization in a continuous pattern along cortical microtubule (Figure 3 a). Protein interaction between SlMAP70-1 and SUN10/SlIQD21a was validated using three different approaches, including Y2H, BiFC and Co-IP (Figure 3 b-d), confirming SUN10/SlIQD21a as bona fide SlMAP70-1 interactor. Such interaction is likely conserved among SlMAP70 homologues, as indicated by Y2H-interaction assays between the five tomato MAP70 isoforms with SUN10/SlIQD21a (Figure 3b, Figure S8 a-c).

**Figure 3.**
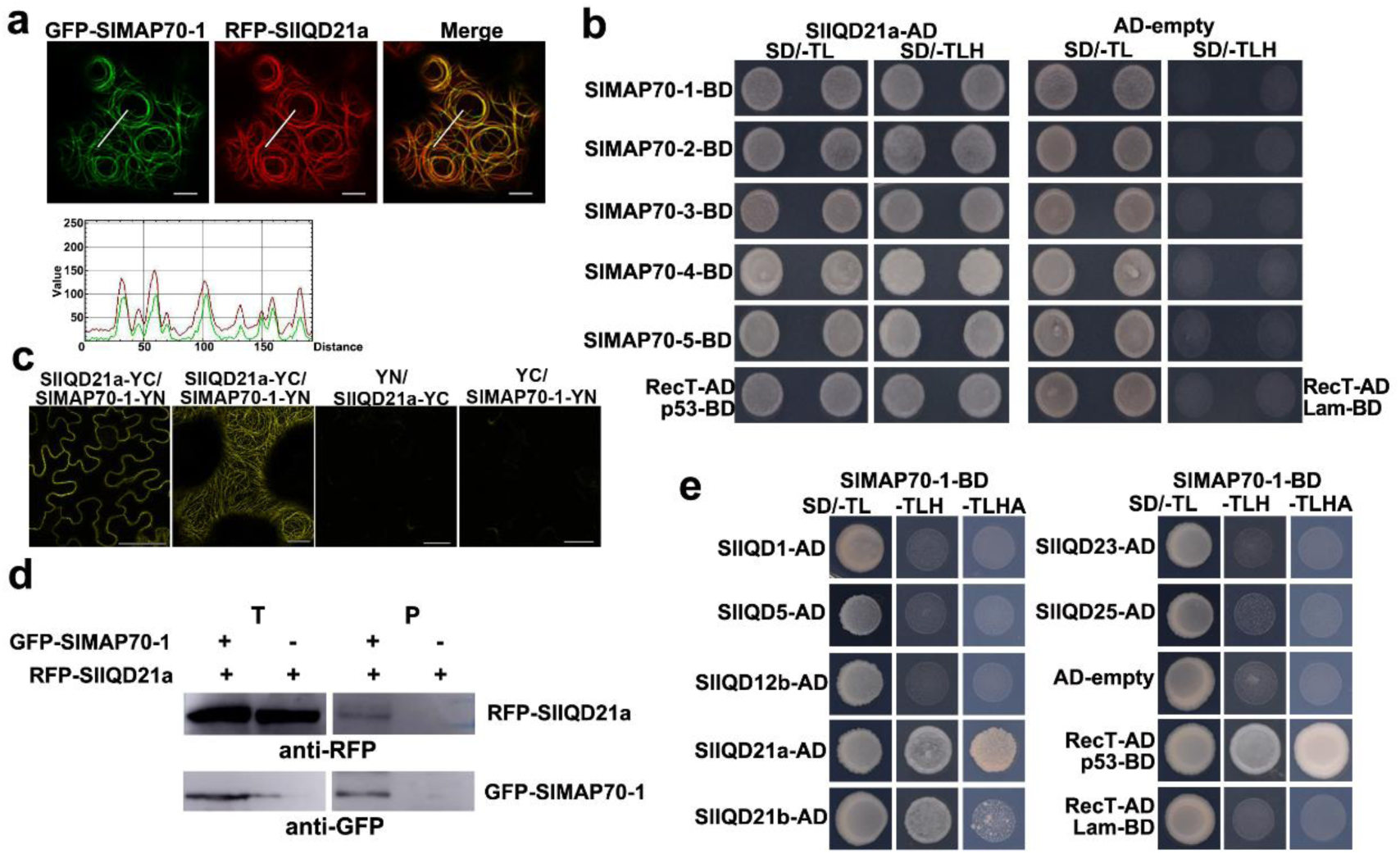
SlMAP70 proteins interact with tomato IQ domain protein 21. **(a)** RFP-SlIQD21a co-localized with GFP-SlMAP70-1 in *N. benthamiana* leaf epidermal cells. The GFP signal of SlMAP70-1 fusion protein along the white line superposed with the RFP signal of SlIQD21a fusion protein. Bars: 10 μm. **(b)** Y2H analysis indicates tomato IQD21a interacts with all tomato MAP70 proteins. p53 + RecT indicates positive control, and Lam + RecT indicates negative control. For Y2H, all the strains could grow on SD/-TL (SD/-Trp/-Leu) media, whereas only positive strains could grow on SD/-LTH (SD/-Trp/-Leu/-His) media, and strong positive strains also could grow on SD/-LTHA (SD/-Trp/-Leu/-His/-Ade). **(c)** BiFC assays of protein interaction between SlMAP70-1 and SlIQD21a in *N. benthamiana* leaf epidermal cells. YFP fluorescence was visible along the MT filamentous structure. Free YN and YC were served as negative controls. Bars: 10 μm. **(d)** GFP-Trap based co-IP assay of GFP-SlMAP70-1 and RFP-SlIQD21a, blot was detected using an RFP antibody. RFP-SlIQD21a was only found in the pellet fraction in the presence of GFP-SlMAP70-1. **(e)** Yeast two hybrid assay of interactions between SlMAP70-1 and other selected SlIQD proteins.

In tomato, 33 SUN/IQD family members were identified, they are divided into three main clades each containing several subclades according to a phylogenetic analysis in comparison to *Arabidopsis IQD* genes (Huang *et al*., 2013). To test if MAP70 interaction is conserved among IQDs, we cloned tomato *SUN/IQD* genes from the distinct phylogenetic subclades, including SUN12/SlIQD12b, the closest homologue to the known fruit shape regulator SUN1/SlIQD12a, and SUN16/SlIQD21b, the closest homologue to SUN10/SlIQD21a. In addition, we selected SUN29/SlIQD1, SUN31/SlIQD5, SUN23/SlIQD23 and SUN32/SlIQD25. Analysis of the subcellular localization of GFP-SlIQD fusion proteins revealed that all of them localize to microtubules as validated by co-localization with RFP-TUA5 (Figure S7b). Interaction of SlMAP70-1 in yeast-two-hybrid assays was observed only for SlSUN10/SlIQD21a, consistent with previous data, and for SUN16/SlIQD21b, the closest homologue of SlIQD21a (Figure 3e). These findings suggest that SlIQD proteins differentially interact with MAP70 proteins, and that interaction may be limited to few IQD proteins.

Previous study reported that the tomato *sun* variant showed a long fruit phenotype, which is attributed to higher expression levels of SUN1/SlIQD12a (Xiao *et al*., 2008). To more broadly test the function of tomato IQDs in fruit formation and shape establishment, we selected three of the GFP-SUN/SlIQDs for further functional analysis. We included SUN10/SlIQD21a, the MAP70 interacting protein, SUN12/SlIQD12b, the homologue of SUN1 (Huang *et al*., 2013; Xiao *et al*., 2008), and SUN29/SlIQD1, which shares sequence similarity with *Arabidopsis* IQD2 involved in root growth regulation (Zang *et al*., 2021). Over-expression constructs of GFP-SUN/SlIQD fused variants were transformed into tomato plants. As in *N. benthamiana*, all three GFP-SUN/SlIQD variants labeled microtubules as demonstrated by oryzalin treatment (Figure S9a). Interestingly, fruits from these transgenic plants showed different shapes. *SUN29/SlIQD1* OE lines displayed no significant changes in fruit size and shape compared to wild type, while the fruits from both *SUN12/SlIQD12b* and *SUN10/SlIQD21a* OE lines were elongated (Figure S3d, Figure 4a, Figure S9 b-c). In *SUN12/SlIQD12b* transgenic plants, fruits were extremely elongated, similar to phenotypes reported for *SUN1/SlIQD12a* overexpression plants (Xiao *et al*., 2008). Fruits from the *SUN10/SlIQD21a* transgenic plants exhibited only mild elongation, similar to those in *SlMAP70-1* OE plants (Figure 1d; Figure 4a). Therefore, we suspected that different SUNs/SlIQDs may control fruit shape via different molecular mechanisms, and that SUN10/SlIQD21a and SlMAP70-1 may have a similar or joint function on microtubule organization, and regulate fruit shape coordinately.

**Figure 4.**
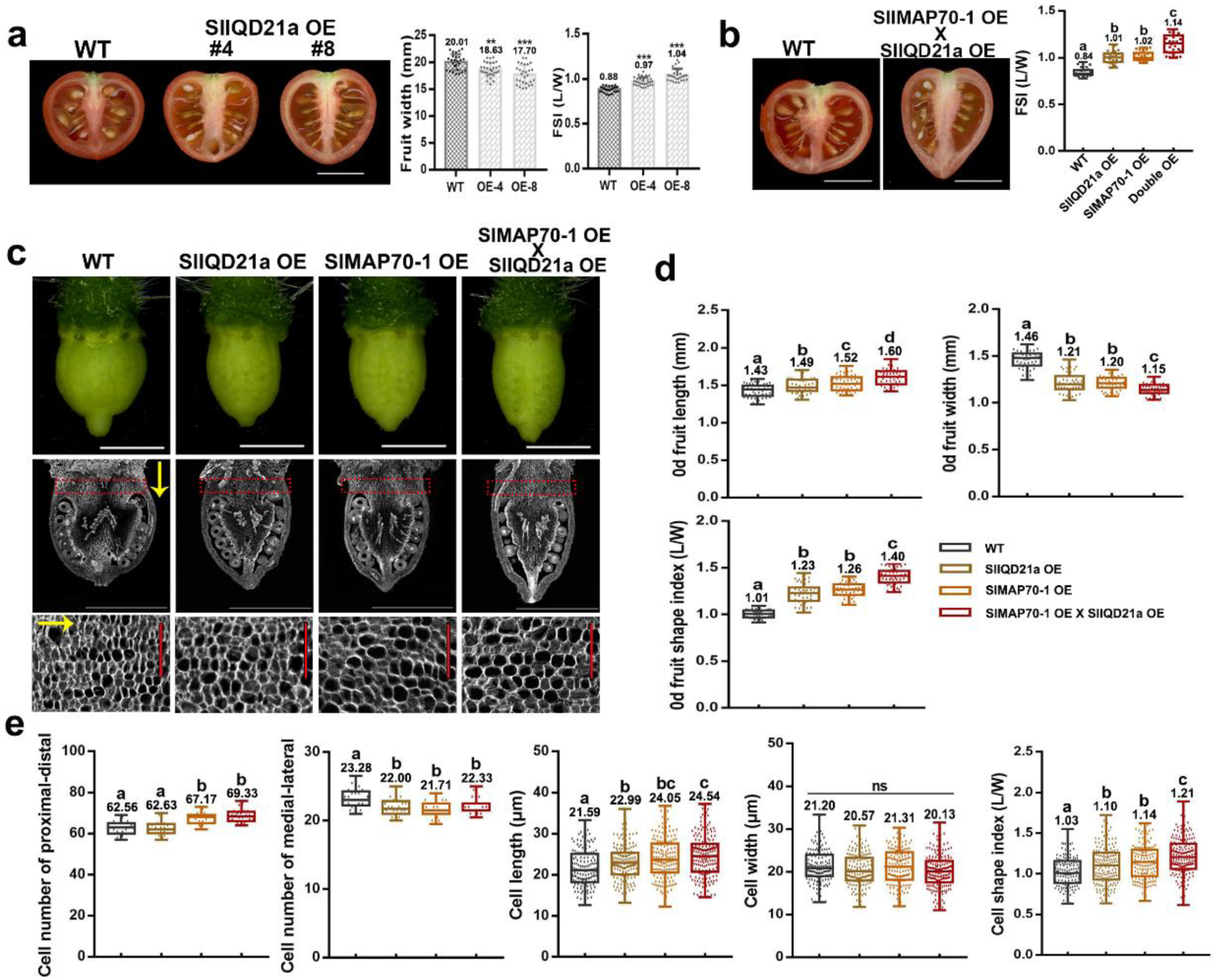
Co-expression of *SlIQD21a* and *SlMAP70-1* further promote tomato fruit elongation. **(a)** Overexpression of SlIQD21a increased fruit shape index. Data are shown as mean ± SD, n=30. ***, P < 0.001; **, P < 0.05 by Tukey’s test. Bars: 1 cm. **(b)** Overexpression of *SlIQD21a* and *SlMAP70-1* together results in a longer fruit phenotype compared to single overexpression. The letters in the graph indicate the significant differences in fruit shape index evaluated by Tukey’s test (p < 0.05), respectively. **(c)** Microscope images of anthesis-stage ovaries (up) and paraffin sections along longitudinal axis (medium). Images at the bottom are cells from the proximal end of ovaries with a high magnification. Yellow arrows indicate the proximal-distal direction of ovary growth. White bars: 1 mm. Red bars: 0.1mm. **(d)** Statistical analysis of fruit length, width and aspect ratio in (c). **(e)** Cell numbers along proximal-distal and medial-lateral direction, cell length, width and aspect ratio in the parenchyma zone (red dotted rectangle) of anthesis ovaries in (c). The letters indicate the significant difference evaluated by Tukey’s test (α < 0.05), respectively.

### Co-expression of SUN10/SlIQD21a and SlMAP70-1 further enhances fruit elongation

To further dissect the proposed joint role of SlMAP70-1 and SUN10/SlIQD21a in fruit shape regulation, we crossed *SlMAP70-1* OE lines with *SUN10/SlIQD21a* OE lines, generating stable tomato plant expressing both constructs. Compared to the *SlMAP70-1* and *SUN10/SlIQD21a* single OE lines, the *SlMAP70-1; SlIQD21a* double OE lines showed an enhanced fruit elongation phenotype and larger fruit shape index. We hypothesize that SlMAP70-1 and SUN10/SlIQD21a have joint effects on fruit elongation (Figure 4b). According to the gene expression data (Figure 1b, S7a), we suspected that both SlMAP70-1 and SUN10/SlIQD21a may exert their function from early fruit set stage on. Therefore, ovaries right after anthesis were studied at both tissue and cell level. When compared with wild-type tomato, all single and double *SUN10/SlIQD21a* and *SlMAP70-1* OE lines displayed significant changes in ovary shape (Figure 4c). *SlMAP70-1* OE lines exhibited a slender ovary with a significant increase in ovary length and decrease in a width, similar to the phenotype of *SUN10/SlIQD21a* lines. Co-overexpression of *SUN10/SlIQD21a* and *SlMAP70-1* further enhanced ovary elongation and reduced ovary width (Figure 4d), in agreement with the shape changes of the mature fruits (Figure 4b).

To reveal the mechanisms that underlie these morphological changes, cell parameters at the proximal end (Figure 4c, red rectangle) of the anthesis-stage ovaries were characterized according to Wu et al (Wu *et al*., 2018). Cell width was barely altered in either *SlMAP70-1* or *SUN10/SlIQD21a* OE line compared to wild type, whereas cell length was significantly longer in all transgenic lines (Figure 4e). Also, cell number of the whole fruit increased significantly along the proximo-distal axis and decreased along the medial-lateral axis in all *SlMAP70-1* OE lines. These results suggested that the main role of SlMAP70-1 and SUN10/SlIQD21a in fruit elongation is to regulate cell expansion at the longitudinal direction, and possibly affect cell division from early ovary stages on.

### SUN10/SlIQD21a and SlMAP70-1 disrupt microtubules dynamic re-arrangement during early fruit development

Microtubule arrangement can be influenced by several factors, such as cell shape and space confinement (Colin et al., 2020). Therefore, we decided to focus on the late ovary development stages (-3 d to 0 d) and early fruit development stages (0 d to 3 d) to study the function of SlMAP70-1 in microtubules dynamical rearrangement. Cells from different fruit tissues were checked (Figure S10a), we found only cells from the endocarp tissue are suitable for subcellular studies of microtubules. In other tissues of the fruit, cells are either very small (e.g exocarp pavement cells) or not easily accessible (e.g placenta and mesocarp cells; Figure S10b). Our results showed that the average cell area of endocarp cells exhibited little variation at this stage, and their cell circularity remained the same (Figure 5a-c), therefore the variation of cell shapes to microtubules structures were minimal. To gain insights into the organization and dynamics of microtubules in endocarp pavement cells during fruit development, we employed a transgenic tomato line expressing the GFP-MAP65-1 (Figure 5 a-b), which was used as a microtubule marker in previous studies (Riglet et al., 2020). We also attempted to generate lines expressing other microtubule markers (e.g., RFP-TUA5), but these fusion proteins have dramatic impact on tomato reproduction and no transgenic seeds were obtained. Prior to our detailed analyses, we validated that GFP-MAP65-1 completely co-localized with the microtubule marker RFP-TUA5, and that expression of GFP-MAP65-1 had no effect on tomato fruit size and shape (Figure S11 a-b), suggesting that GFP-MAP65-1 is suitable to be used as a marker for analysis of cortical microtubules in tomato fruits. Analysis of GFP-MAP65-1 lines revealed that during the early fruit development the value of microtubule anisotropy decreased gradually from days -3 to day 0 (the day of anthesis), meaning microtubules became more disordered in this period. From day 0 to day 3, microtubule anisotropy increased significantly to levels higher than the -3d stage, and dropped again from day 3 afterwards (Figure 5 c). These changes in microtubule organization indicate a precisely regulated machinery controls the dynamic re-orientation of microtubule during progression of fruit development, in which MAPs are likely be involved.

**Figure 5.**
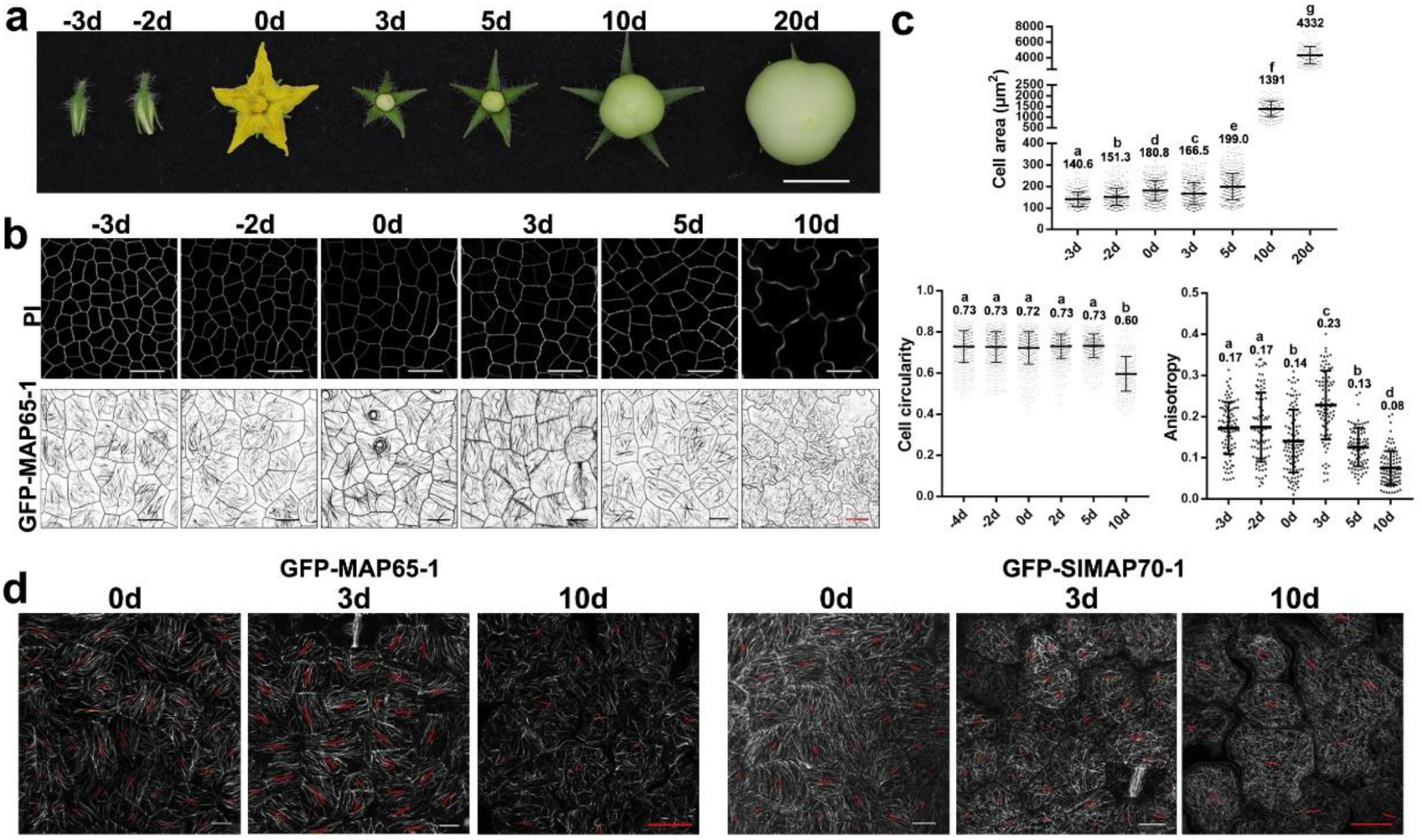
SlMAP70 and SlIQD21a regulate cortical microtubule arrangement in tomato endocarp cell during fruit growth. **(a)** Developing tomato fruits at -3, -2, 0, 3, 5, 10 and 20 days after anthesis. **(b)** Endocarp pavement cells at different fruit development stage, and its microtubules arrays labeled by GFP-MAP65-1. Black bars: 10μm. White or red bars: 25μm. **(c)** Quantitative analyses of endocarp pavement cell size, circularity, and microtubules anisotropy in wild-type endocarp cells of different fruit development stages. At early developmental stages (day 0), cells exhibited random orientation of microtubules arrays, it became increasingly ordered into parallel linear orientation at active stage of cell division (day 3), then return to random orientation at cell enlargement stages (10 days after anthesis). The letters indicated a significant difference by Tukey’s multiple comparisons test (α<0.05). Values are given as the mean ± SD of about 100 cells. **(d)** Image of CMTs in tomato endocarp cells of MAP65-1 and MAP70-1 transgenic lines. The length and orientation of red lines represent the anisotropy and average orientation of the CMT arrays that quantified with FibrilTool. Bars: 10 μm.

In wild type tomato endocarp cells, GFP-MAP65-1 labelled microtubules exhibited a highly disordered arrangement on the day of anthesis (day 0), and became more parallel-arranged/ordered 3 days later when cell division was active (Figure 5 d). In *GFP-SlMAP70-1* over-expression lines, the microtubules appeared to be more parallel at day 0, while became less ordered at day 3 (Figure 5d), such behavior appeared very different from the microtubule organization observed in wild type tomato at the same stage (Figure 5c). The length of each microtubule filament also appeared to be longer in *SlMAP70-1* OE lines, especially during fruit enlargement stages (day 10) (Figure 5d). Taken together, our results indicated that microtubules rearranged dramatically during early fruit development, and this is likely important for cell and fruit morphology. The precise regulation of microtubule patterning could be mediated by various microtubule binding proteins, such as SlMAP70 that exhibited elevated expression in young fruits.

### SUN10/SlIQD21a and SlMAP70 proteins affect microtubule stability

So far, we showed that the expression level of SlMAP70, and its interaction with SUN10/SlIQD21a is essential for fruit elongation. Next, we moved on to study the exact functions of SlMAP70 on microtubules using chemical treatment and image analytical methods. In *N. benthamiana* cells, co-expressing RFP-SUN10/SlIQD21a and GFP-SlMAP70 induced the formation of thick microtubule bundles and microtubule-whorls were predominant (Figure 3a, Figure 6a, Figure S8c). Similar microtubule pattern had been reported in a few previous studied, but their biological relevance is not clear (Kirik et al., 2007; Ren et al., 2017). From a biophysical perspective, it is suggested that microtubule-whorls are promoted by local interactions between each filament (e.g proteins induced microtubule bundling), and results from a joint effect of microtubule collision followed by reptation-like motion of single microtubules (Sumino et al., 2012). To gain quantitative insights into the changes in microtubule organization, we measured the length of the microtubule filaments. Extreme long filaments (>50 µm) were very abundant in cell expressing *SlMAP70-1* or both *SlMAP70-1* and *SUN10/SlIQD21a*. This abnormal behavior may in favor of formation of microtubule-whorls, which consisted of very long microtubule filaments, in either clockwise or anti-clockwise direction (Figure 6b).

**Figure 6.**
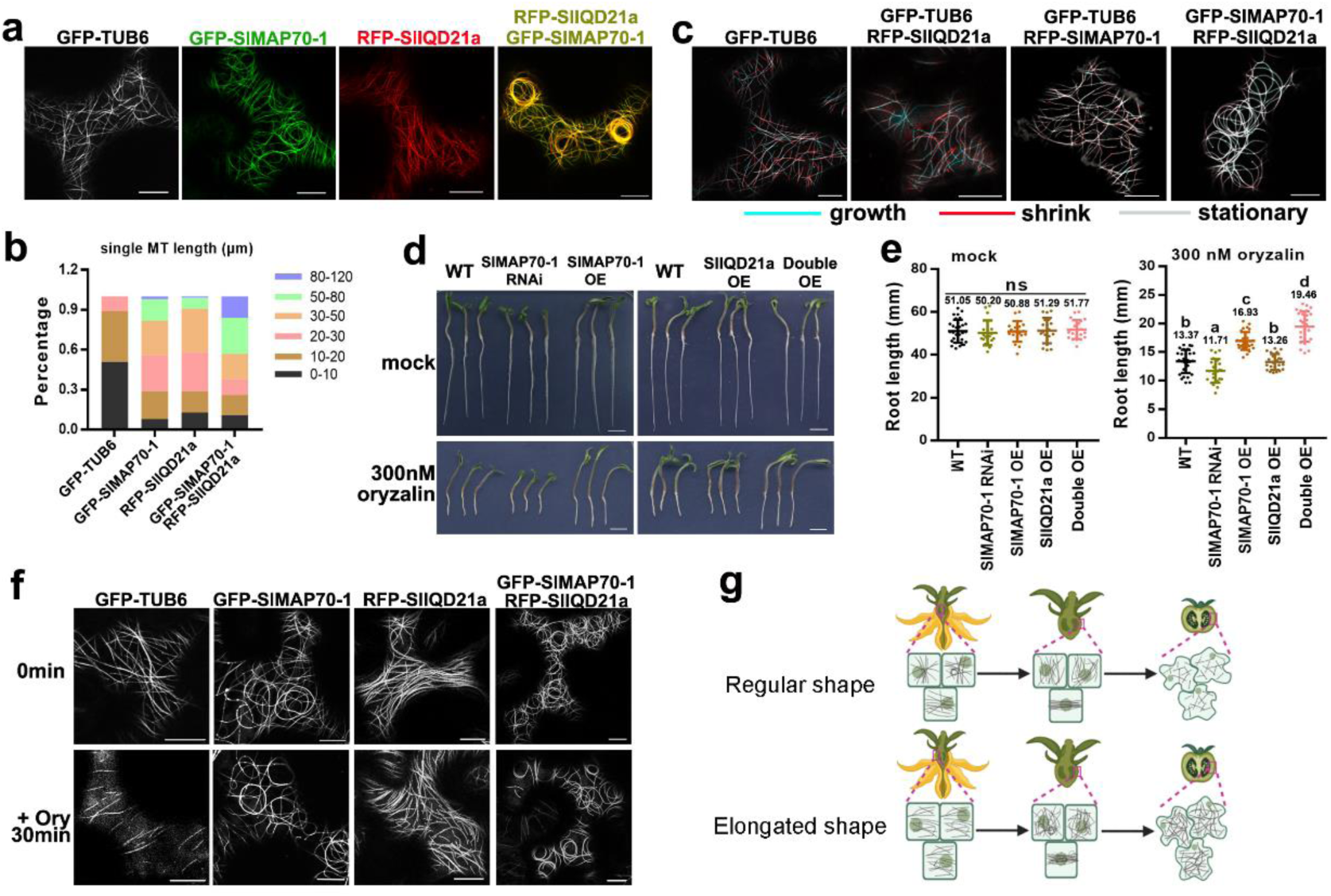
Over-expression of SlMAP70-1 and SlIQD21a proteins stabilize microtubules *in vivo*. **(a)** Co-expression of RFP-SlIQD21a and GFP-SlMAP70-1 resulting in long and bundling microtubules that prone to form whorls-like array in *N. benthamiana* leaf epidermal cells. Bars: 10 μm. **(b)** Statistical analysis of microtubule filament length in (a). The colors indicate the different length ranges of microtubule microfilaments. More than 100 filaments were analyzed. **(c)** Representative confocal images from a time lapse in *N. benthamiana* leaf pavement cells showing dynamic microtubules in color and preexisting microtubules in gray. Green line indicates newly grown cMT, and red line indicates shrink cMT, respectively. A slight growth and shrinkage of cMTs were detected with the co-expression of GFP-SlMAP70-1 and RFP-SlIQD21a proteins. A total of ten repeats were performed for each construct. The frames are separated by 45 s. Bars: 10 μm. **(d)** Oryzalin treatment of different SlMAP70-1 genotypes. Germinated seedlings were grown on regular medium supplemented with 0 or 300 nM oryzalin for 3 days. Bars: 1cm. **(e)** Statistical analysis of root lengths of indicated genotypes under oryzalin treatment in (d). The *SlMAP70-1; SlIQD21a* co-overexpression lines were the most resistant to oryzalin treatment. Different letters in dot plots indicate the significance. ns: no significant, p < 0.05 by Tukey’s test. n = 25. **(f)** Effect of oryzalin on microtubules in *N. benthamiana* leaf pavement cells expressing GFP-TUB6, or GFP-SlIQD21a and GFP-SlMAP70-1. Representative confocal images were captured before and after treating with 100 μM oryzalin for 30 min, respectively. Bars, 10 μm. **(g)** A schematic model of tomato fruit shape establishment.

To determine the effect of SUN10/SlIQD21a and SlMAP70-1 expression on microtubule dynamics, time-lapse studies were performed using a frame-subtraction approach (Lindeboom et al., 2013; Smertenko et al., 2020). Here, microtubule growth (polymerization) was pseudocolored in red and microtubule shrinkage (depolymerization) were pseudocolored in cyan. In control cells expressing GFP-TUB6, both events were frequently observed, while microtubule dynamic appeared to be reduced in the presence of SUN10/SlIQD21a or SlMAP70-1. In cells co-expressing *SUN10/SlIQD21a* and *SlMAP70-1*, microtubules appeared static as little growth or shrinkage were observed in the processed images (Figure 6c; Movie S1, S2). Thus, these results suggested that SUN10/SlIQD21a and SlMAP70-1 are able to stabilize microtubules, and that this effect was further enhanced when both proteins are co-expressed.

To confirm the possible role of SlMAP70-1 and SUN10/SlIQD21a in regulating the stability of microtubules, tomato seedlings were treated with 300 nM oryzalin, which prevents de novo microtubule polymerization and inhibit plant growth (Morejohn *et al*., 1987; Yang *et al*., 2020) (Figure 6d). Root growth without oryzalin treatment was similar among different genotypes. However, tomato seedlings over-expressing *SlMAP70-1*, or *SlMAP70-1*; *SUN10/SlIQD21a* were more resistant to oryzalin treatment, exhibiting longer roots after oryzalin treatment, while plants with reduced expression of *SlMAP70s* were more susceptible (Figure 6e). High concentration oryzalin treatment disrupted most microtubule that labeled by GFP-β-Tubulin6 (GFP-TUB6) within 30 min (Figure 6f), but this effect was less prominent in the presence of over-expressed *SlMAP70-1* or *SUN10/SlIQD21a*. All these results support our conclusion that SlMAP70 likely regulate microtubule function through a stabilizing effect, which was further enhanced when *SlMAP70-1* and *SUN10/SlIQD21a* are co-expressed (Figure 6f). Such stabilization effect may perturb microtubule dynamical rearrangement in various cell types (Figure 2, Figure 5), and directly attributed to the alternation in cell directional growth and fruit shape establishment (Figure 6g).

### Conclusion and perspective

During the past years, genes encoding microtubule binding proteins emerged as conserved trait loci for fruit shape regulation with important roles during domestication, as has been reported in tomato, cucumber, melon, peach and rice (Dou *et al*., 2018; Pan *et al*., 2020; Wu *et al*., 2018; Yang *et al*., 2020; Zhou et al., 2021). However, most of these studies were performed using forward genetics or population genetics approach, the in-depth molecular and cell biological insight have not been investigated. Early studies in model plants showed that MAPs regulate microtubule function in different ways, contribute to cell wall composition, and cell polarity and cell division, understanding the molecular basis of these processes in fruit crops would certainly provide a powerful toolbox for fruit shape regulation in the future. Here, we have for the first time studied the dynamics of microtubules in tomato fruits, and identified novel candidates for fruit shape formation. Over-expressing *SlMAP70-1* promotes microtubule stabilization and reduces cytoskeletal dynamics that correlate with the formation of elongated fruits. The loss-of-function of *SlMAP70* has the opposite effect confirming that SlMAP70s are important regulators of tomato fruit shape. We further demonstrate that SlMAP70 interacts with members of the SUN/IQD family, in particular with SUN10/SlIQD21a and related proteins, thereby linking functions of these two plant-specific families of MAPs. Both *MAP70-1* and *SUN10/SlIQD21a* genes exhibit high level of expression in young fruits, and may regulate fruit shape formation jointly from early stages of anthesis on. Further studies on microtubule functional regulation may provide a powerful approach for fruit shape regulation in fruit crops.

## Materials and methods

### Gene cloning and plasmid construction

To generate over-expression transgenic plants, the CDs sequences of *SlMAP70-1* (Solyc05g017980), *SUN10/SlIQD21a* (Solyc03g083100) and *SlMAP65-1* (Solyc07g064970) were cloned into pMDC43 to generate N-terminal GFP fusions via Gateway LR ligation. To generate RFP fusion of microtubule marker, the CDS of *Arabidopsis TUA5* (AT5G19780) was cloned into pK7WGR2. For the RNAi construct of *SlMAP70*, a gene-specific fragment (472-bp) of *SlMAP70-1* (Figure S2a) was cloned into the pHellsgate8 vector using LR Clonase (Invitrogen). For the CRISPR/Cas9 constructs of *slmap70*, the target sequences of SlMAP70-1, SlMAP70-2, SlMAP70-4 and SlMAP70-5 were designed using the online CRISPR-P2.0 web tool (http://crispr.hzau.edu.cn/CRISPR2/) and fused with two single-guide RNA (sgRNA) expression cassettes into pTX vector (Figure S3). For making fluorescence protein fusions of SlIQDs, full length sequences were cloned into various destination vectors of pK7WGR2 (N-terminal RFP), pMDC43-GFP (N-terminal GFP) and pK7WGC2 (N-terminal CFP) according to the requirement. The primers used for cloning are listed in *SI Appendix*, Table S1. All constructs were transformed into *Agrobacterium tumefaciens* (strain GV3101) for transformation.

### Plant material and growth conditions

Tomato (*Solanum lycopersicum*) cv. MicroTom was used as the wild type in this study. Tomato transformation was performed as described previously (Sharma et al., 2009). Homozygous lines were used for subsequent analyses. The *Pro35S:GFP-SlMAP70-1* and *Pro35S:GFP-SlIQD21a* co-expressing plants were generated by crossing of single stable lines. The transgenic plants of *N. benthamiana* were generated following a published protocol (Clemente, 2006). All the tomato and *N. benthamiana* plants were grown under a 16-h/8-h light/dark photoperiod in a growth room at 25 °C.

### Fruit and ovary shape analysis

Mature tomato fruits were cut longitudinally and scanned at 600 dpi. Anthesis ovaries were imaged using the Leica M205 FA dissecting scope. The maximum length and width were measured using Image J software, and the Fruit Shape Index was calculated. Three to five plants of each genotype were analyzed, and the average values were taken from at least 25 fruits or ovaries and analyzed with Tukey’s test (α < 0.05).

### Tissue fixation and sectioning

Anthesis ovaries were cut longitudinally with a razor blade and fixed in PFA (4% Paraformaldehyde, 0.01M PBS buffer, pH 7.4) at 4℃ overnight. The samples were dehydrated with ethanol-ddH_2_O series (50, 70, 85, 95, 100%, 100%) at one hour for each step. Then the samples were treated with 1:1 ethanol: poly distearate (Sigma-Aldrich) for 2 h followed by 100% poly distearate at 38°C for 3h. Samples were then solidified in poly-distearate following the paraffin section’s protocol. Embedded samples were cut into 12-μm sections using a Leica RM2255 automated microtome. The sections were stained with 0.01% calcofluor white for 2 min, washed with water and then imaged using a Leica TCS SP8 confocal microscope. Cell size and number were analyzed using the Image J. The length and width of parenchyma cells in the proximal area were evaluated on at least 30 cells per sample. Numbers of parenchyma cells were counted in the columella area in the proximo-distal and the proximal area in the medio-lateral direction.

### Confocal microscopy and image analysis

*N. benthamiana* leaves were pressure-infiltrated through the abaxial epidermis with *Agrobacterium tumefaciens* GV3101 cells containing the construct and the silencing suppressor p19 in a 1:1 ratio. To analysis the subcellular localization of SlMAP70 proteins during cell division cycle, the Pro35S: AtCYCD3;1 construct was also co-infiltrated as described (Xu *et al*., 2019). The images were obtained two days after infiltration. Images were taken with a LEICA TCS SP8 inverted microscope using a 63 x oil-immersion objective. The excitation wavelengths for calcofluor white, GFP, YFP, RFP/PI were 405 nm, 488 nm, 514 nm and 561 nm, respectively, and emission was detected between 410 and 450 nm (calcofluor white), 495 and 550 nm (GFP), 520 and 550 nm (YFP), 590 and 640 nm (RFP/PI). Fluorescence intensity profiles of GFP and RFP fluorescence were generated using the Plot Profile module of Image J. Time lapses of microtubule dynamics were acquired at 15 s per frame. Microtubule ends were artificially color-labelled using a published procedure using ImageJ (Lindeboom *et al*., 2013; Smertenko *et al*., 2020).

To visualize the organization of cortical microtubules in fruit cells, the endocarp pavement cells of ovary wall or pericarp tissues from different fruit development stages were carefully dissected from the middle of the fruits (Figure S9c). Then the samples were put onto a micro slide and the endocarp faced upward for observation. A cover slide was put on the samples gently, and the images were taken using confocal with settings for GFP fluorescence. For quantification of cortical microtubules arrangement in the leaf and endocarp pavement cells, the anisotropy was analyzed by the Fibril Tool plugin of ImageJ (Boudaoud *et al*., 2014). For quantification of cell shape, samples were immerged into a staining solution containing 10 μg/ml propidium iodide (Sigma-Aldrich) and incubated for at least 10 min. Leaf and endocarp pavement cells were imaged by a confocal microscope with RFP setting, and quantified by the PaCeQuant plugin of Image J (Möller et al., 2017).

### Bimolecular fluorescence complementation (BiFC) assays

The CDs of *SlIQD21a* were cloned into p35S:YFPC, and CDs of *SlMAP70-1*, *SlMAP70-2* and *SlMAP70-3* were cloned into p35S:YFPN (Jakobson et al., 2016) vector by gateway LR reaction respectively. Resultant constructs and empty vectors were co-expressed in *N. benthamiana* leaves and YFP fluorescence was observed by Leica TCS SP8 confocal microscope after infiltration for 3 days.

### Yeast-two-hybrid (Y2H) studies

Full-length of *SlMAP70s* were cloned into the pGBKT7 vector, and full-length *SlIQDs* selected were cloned into the pGADT7 vector by gateway LR reaction. A pair of bait and prey constructs were co-transformed into the AH109 yeast strain and spotted on vector-selective medium lacking Trp and Leu (SD/-Trp/-Leu). The yeast transformants were screened on interaction-selective medium lacking Trp, Leu and His (SD/-Trp/- Leu/-His).

### Antibody production

DNA fragments corresponding to antigen peptide of SlMAP70-1 (amino acid residues 397 to 501) were cloned into pET28a vector. The antigen peptide was expressed with a 6xHis tag fusion on the N terminus (pET28a) in E. coli BL21+, then purified according to the instruction of Ni-NTA Agarose (70666-4, EMD Millipore Corporation). Purified proteins were used to raise antibodies in mice as described in Smertenko, et al (Smertenko et al., 2008). The polyclonal antibodies were tested by western blot in tomato and *N. benthamiana* plants as shown in Figure S6.

### Western blot and coimmunoprecipitation (Co-IP) assay

Samples of young leaf were collected from one-month old seedlings of individual genotypes, tissues were grinded in liquid nitrogen, and resuspended in 2×SDS buffer (0.125 M Tris-HCl pH 6.8, 200mM DTT, 4% SDS, 20% glycerol, 0.2% bromophenol blue) for crude protein extraction. The anti-GFP antibody (Biorbyt, 1:1000 dilution, mouse), anti-MAP70 antibody (1:500 dilution, mouse), anti-RFP antibody (Abcam, 1:1000 dilution, rabbit) and anti-actin antibody (BBI, 1:1000 dilution, rabbit) were used as the primary antibody (25℃, 2hr). And the Goat anti-mouse (Yeasen, 1:5000 dilution) and Goat anti-rabbit (BBI, 1:5000 dilution) were used as the secondary antibody (25℃, 1hr) according to the host species of first antibody. For detection, the membrane was incubated in enhanced chemiluminescence (ECL) solution (36222ES60, Yeasen) and imaged using an imaging system (FluorChem M, Alpha Innotech).

For GFP-trap based Co-IP assays, GFP-SlMAP70-1 and RFP-SlIQD21a proteins were co-expressed in *N. benthamiana* leaf cells, and single expression of RFP-SlIQD21a was used as a control. Two days after infiltration, total proteins were extracted with lysis buffer (10 mM Tris-Cl pH 7.5, 150 mM NaCl, 0.5 mM EDTA, 0.5% NP-40, 100 mM PMSF, and 1X Roche protease inhibitor cocktail), followed by incubated with anti-GFP beads (AB_2631358, Chromotek) for 2 h at 4°C. The beads were washed at least three times with wash buffer before SDS-PAGE. The immunoprecipitated proteins were analyzed using western blot with anti-GFP antibody (orb323045, Biorbyt) and anti-RFP antibody (ab62341, Abcam).

### Immunofluorescence of tomato leaf cells

The second leaves from 3 weeks old tomato plants were fixed in 4% (w/v) PFA and 0.1% (v/v) Triton X-100 in PEM buffer (50mM PIPES, 5mM EGTA, 5mM MgSO_4_, pH 7.0) for 90 min. After five wash steps with PEM buffer, the samples were frozen, shattered and permeabilized as described (Celler et al., 2016). Then, the samples were blocked in PBS buffer with 2% (w/v) BSA for 2hr and incubated with a primary anti-a-tubulin antibody (AF7010, Affinity, 1:200 dilution) overnight at 4℃, and a secondary antibody (TRITC-conjugated anti-Rabbit IgG, 1:200 dilution, Jackson Immuno Research) for 2hr at 25℃. Washed three times after incubation, the samples were observed using Leica TCS SP8 Laser Scanning Confocal with the RFP settings.

### Oryzalin treatment

For low concentration oryzalin treatment for prolonged period, seeds of different genotypes were grown on 1/2 MS medium. After 1 week, seedlings with similar root length were transferred into new medium supplement with DMSO or 300 nM oryzalin for 4 days, respectively. More than 25 individual roots were taken from every genotype. For high concentration oryzalin treatment, leaf segments of infiltrated *N. benthamiana* expressing various microtubule binding proteins were submerged in 100μM oryzalin solution for 30 min before imaging.

### Phylogenetic analysis and RNA-Seq atlas analysis

To infer the phylogenetic relationships, amino acid sequences of 5 *Arabidopsis* MAP70 proteins and 5 tomato MAP70 proteins were aligned by ClustalW. The phylogenetic tree was generated by the Maximum Likelihood method, using JTT matrix-based model with 1000 rapid bootstrap replications (Kumar et al., 2016). To acquire the tissue-specific transcription data, the ID numbers of five *SlMAP70* genes and 31 *SlIQD* genes were entered to the tomato functional genomics database (http://ted.bti.cornell.edu/). And the reads per kilobase per million reads (RPKM) values were downloaded from the RNA-seq data of tomato cultivar Heinz. The heat map was created using TBtools (Chen et al., 2020).

## Funding

The project was supported by a NSFC grant (no. 91854102) and the Foundation of Hubei Hongshan Laboratory (2021hszd016) to P.W., and a grant (2021CFB142) to Z.B.

## Author Contribution

Z.B. performed the experiments and wrote the manuscript with P.W.; Y.G performed the Y2H related studies; P.W conceived and supervised the project; Y.D., J.Z. and J.Z help on generating transgenic plants and tissue culture; X.Q. helped on confocal microscopy and participated in project progress meetings; K.B. helped on the IQD related studies and imaging analysis; K.B. and B.O. proofread the manuscript and provide critical advice throughout the study.

## Acknowledgements

We thank the support from microscopy core facilities of the College of Horticulture & Forestry sciences, Huazhong Agricultural University. We thank Yuchen Long (National University of Singapore) and Feng Zhao (Northwestern Polytechnical University) for their advice on this work.

## Supplementary Figures

**Figure S1.**
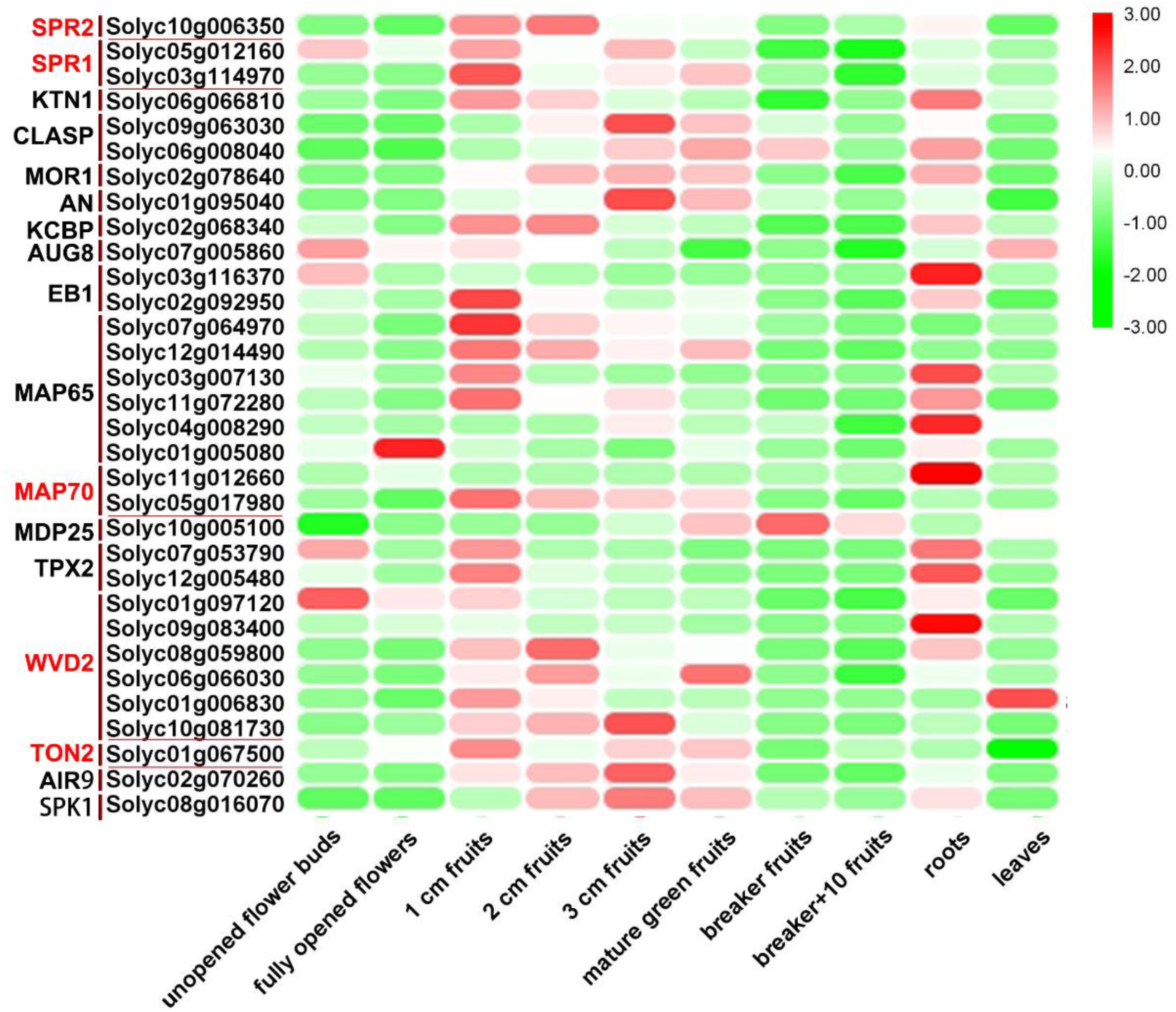
Hierarchical clustering of expression values of some *MAPs* genes in different tissues of Heinz cv. tomato. Red indicates high expression level; green, low expression level.

**Figure S2.**
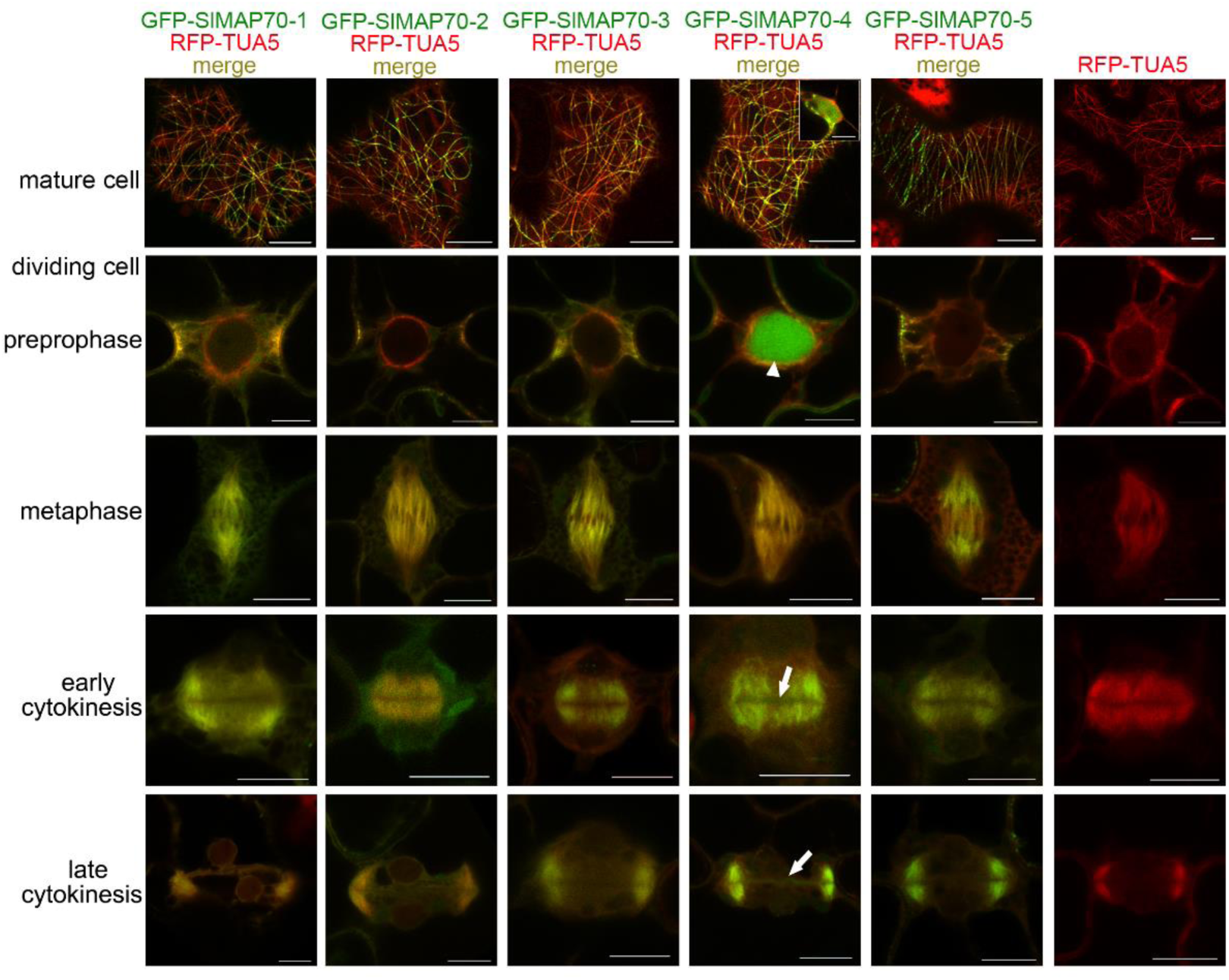
Sub-cellular localization of SlMAP70 proteins. All GFP-SlMAP70 proteins label cortical microtubules, preprophase band, spindle and phragmoplast structures during cell division process in *N. benthamiana*. Whereas, GFP-SlMAP70-4 also label nucleus (arrowheads) in preprophase and cell plate (arrow) at cytokinesis stage. Bars: 10 μm.

**Figure S3.**
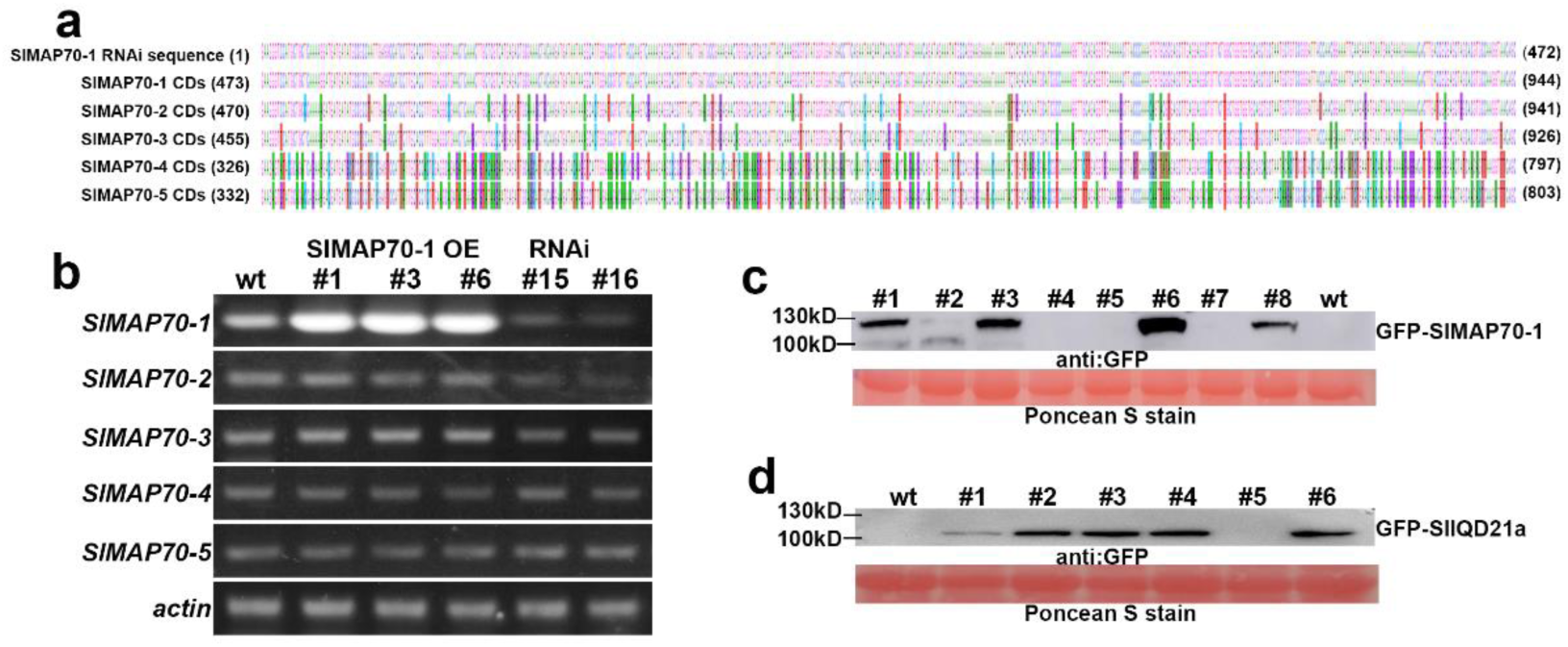
Characterization of *SlMAP70-1* OE and RNAi transgenic plants. **(a)** The alignment of nucleotide sequence of *SlMAP70* genes, in comparison to *MAP70-1* RNAi target sequence. The color box coded nucleotide indicates the difference. The RNAi sequence has a high sequence similarity to *SlMAP70-1* (100% similarity), *SlMAP70-2* (92.05%) and *SlMAP70-3* (90.89%), a lower sequence similarity to *SlMAP70-4* (64.83%) and *SlMAP70-5* (65.25%). **(b)** qPCR analysis of the transcript levels of *SlMAP70* family genes in young leaves of individual genotype. And the expression of *SlMAP70-1*, *-2*, *-3* were all decreased in *SlMAP70-1* RNAi line #15 and #16. *SlMAP70-4* and *-5* has no changes. **(c)** Western blot analysis of the expression level of SlMAP70-1 protein in transgenic lines. SlMAP70-1 highly expressed in line #1, #3, #6, and #8. Ponceau S staining was used as a loading control, and ‘micro tom’ wild-type line as a negative control. **(d)** Western blot analysis of the expression level of SlIQD21a protein in transgenic lines. SlIQD21a highly expressed in line #2, #3, #4, and #6. In both (c) and (d), proteins were detected using a GFP antibody.

**Figure S4.**
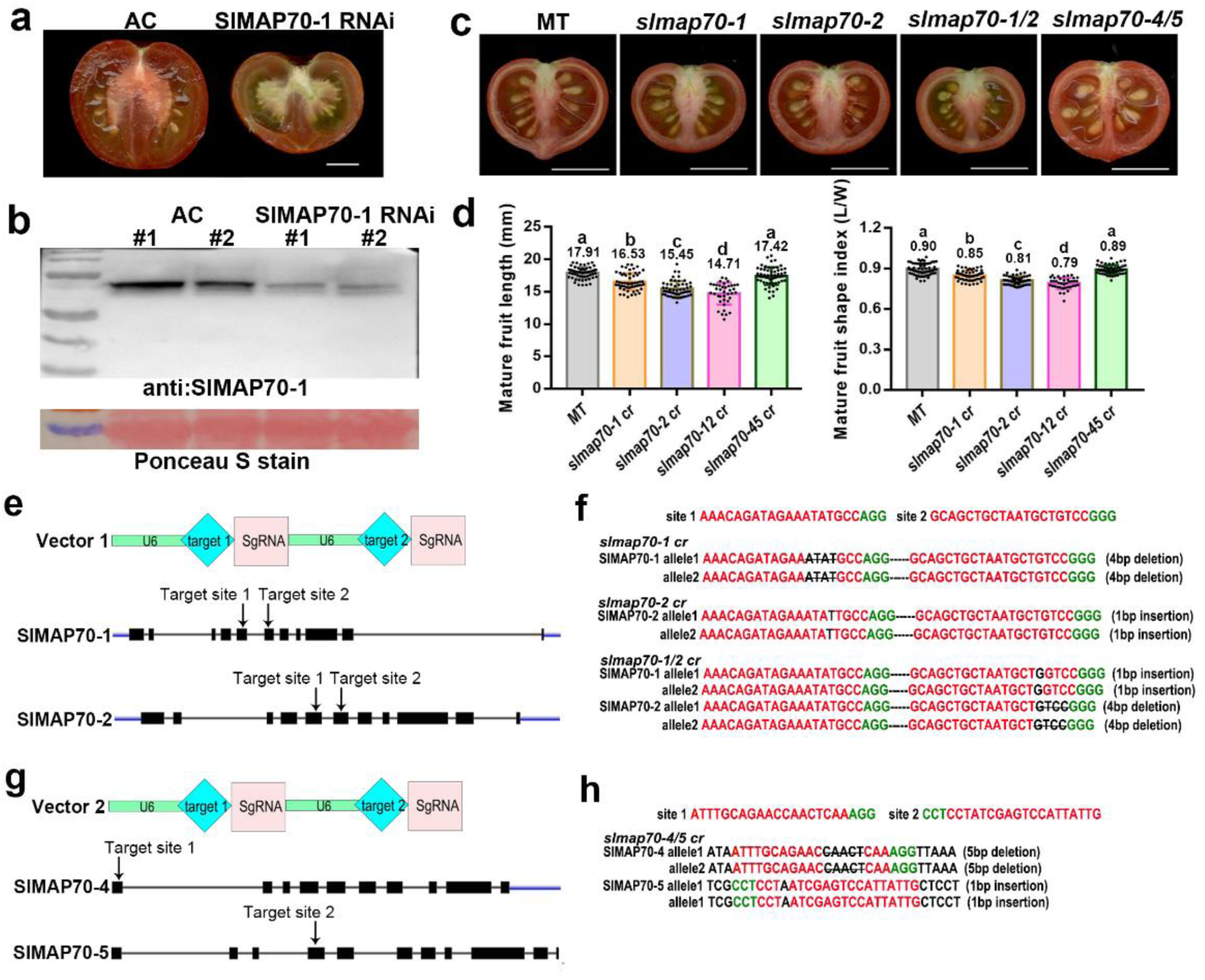
Characterization and phenotype study of *SlMAP70-1* RNAi and *SlMAP70* CRISPR mutant plants. **(a)** Tomatoes of the AC background were transformed with *SlMAP70-1* RNAi construct, and the transgenic line display flat fruit phenotypes. Bars: 1cm. **(b)** The endogenous SlMAP70 protein was reduced in *SlMAP70-1* RNAi lines, as indicated by a western blot using SlMAP70-1 antibody. Ponceau S staining was used as a loading control. **(c-d)** Fruit shape of *SlMAP70* knockout mutant by crispr-cas9. The loss-of-function of *SlMAP70-1* and *SlMAP70-2* result in a flatter fruit, which is similar to the *SlMAP70-1* RNAi line, and the fruit shape of *slmap70-4/5* double mutant has no significant changes compared to ‘MicroTom’ wild-type line. The uppercase letters indicate the difference of p<0.05 by Tukey’s test. Bars: 1 cm. **(e)** The *map70-1/2* single or double mutant were generated by CRISPR/Cas9. In pTX vector 1, two CRISPR/Cas9 target sites were designed to recognize the relative sequences of *SlMAP70-1* and *SlMAP70-2* gene. The arrows indicate the relative location of target sites on *SlMAP70-1* and *SlMAP70-2* gene structure. **(f)** The mutated sites of tomato *slmap70-1/2* single mutant and *slmap70-1/2* double mutant line. **(g)** *The map70-4/5* double mutants were generated by CRISPR/Cas9. In pTX vector 2, the target site 4 could recognize the relative sequence in first exon of *SlMAP70-4* gene, target site 5 recognize the relative sequence in fourth exon of *SlMAP70-5* gene. (**h**) The mutated sites of *tomato slmap70-4/5* double mutant line.

**Figure S5.**
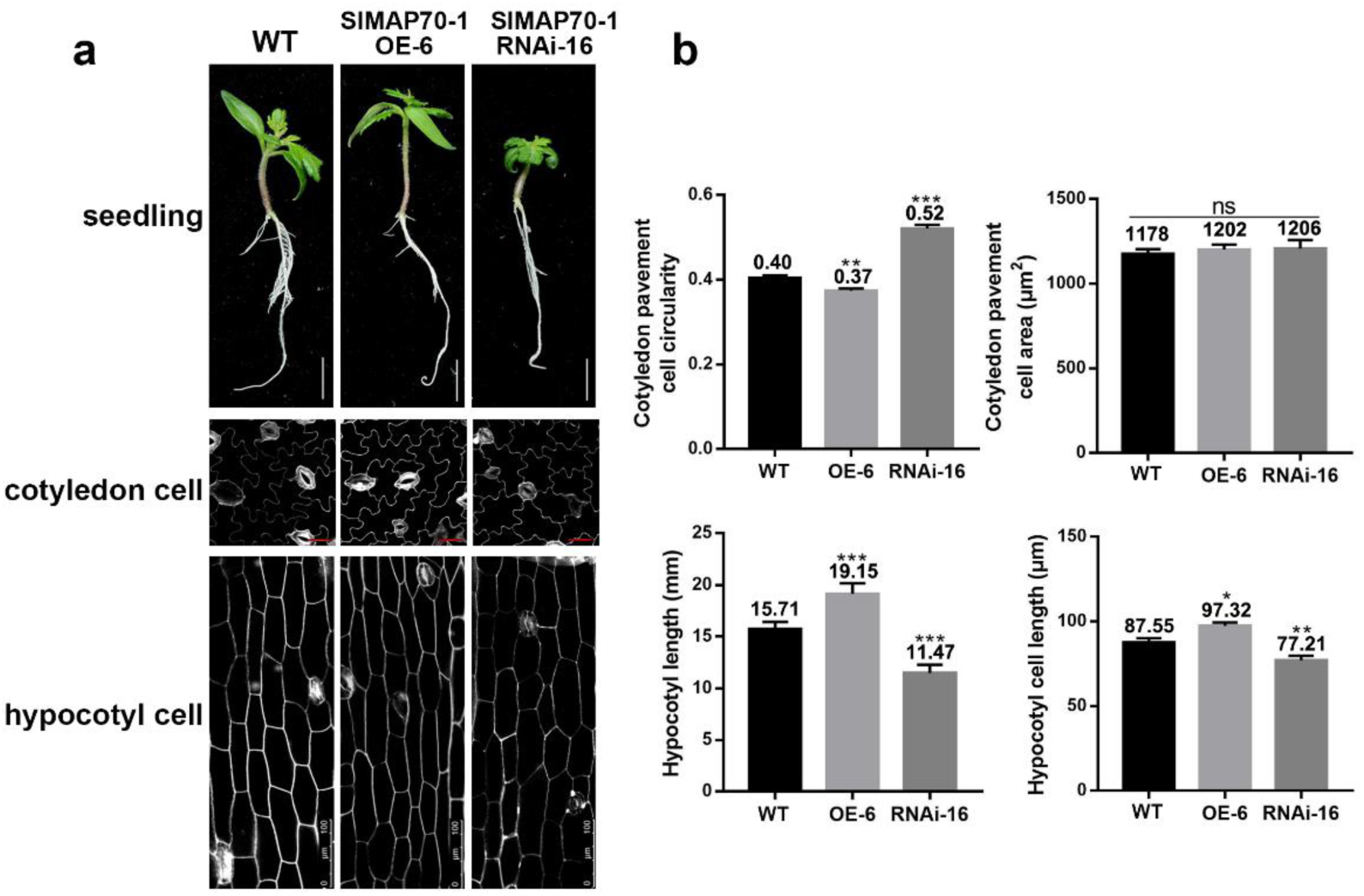
In non-fruit tissue, cell morphology of SlMAP70-1 transgenic plants was also altered. **(a)** Seedling morphology, cotyledon abaxial cells and hypocotyl cells of 15-d-old wild type (WT), *SlMAP70-1* overexpression (OE-6) and *SlMAP70-1* RNAi knockdown (RNAi-16) seedlings. Bars: 1cm for seedlings, 25 μm for cotyledon PCs, 100 μm for hypocotyl PCs. **(b)** Quantification of the cotyledon cell shape and size, hypocotyl length in the genotypes shown in (a). Error bars indicate SD (n > 100 for cotyledon PCs, n > 50 for hypocotyl cells, n = 15 for seedling). The asterisk represents a significant difference between the wild type and SlMAP70-1 OE, RNAi lines (*, p<0.05; **, p < 0.01; ***, p < 0.001, by Turkey’s test).

**Figure S6.**
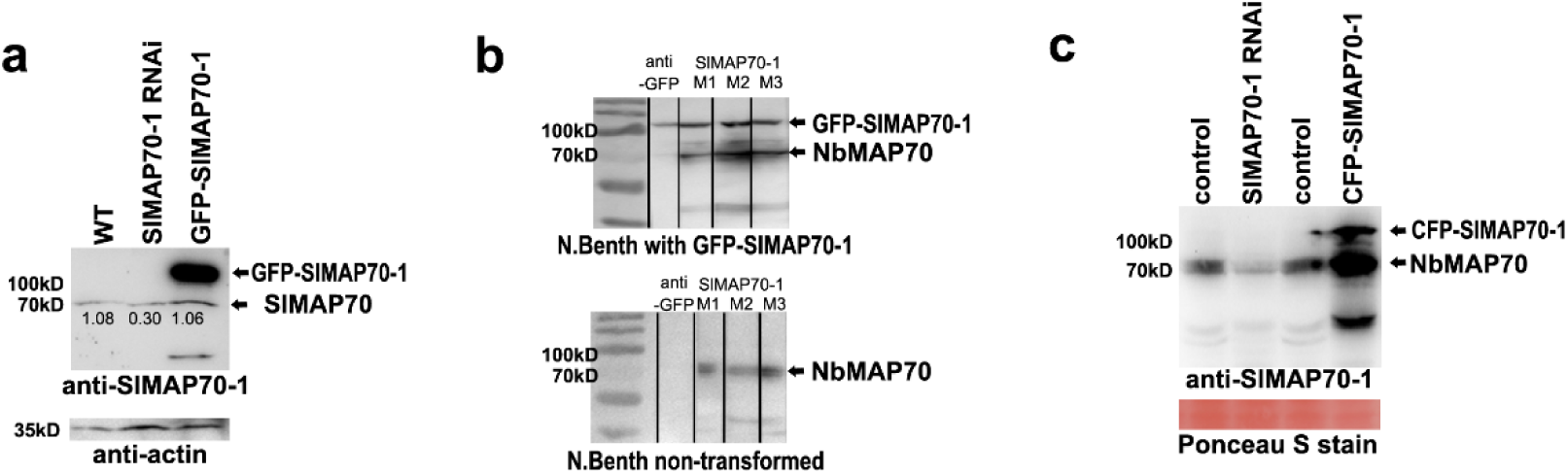
Specificity detection of SlMAP70-1 antibody in tomato and *N. benthamiana* plant. **(a)** Western blot of protein extract from wildtype and transgenic tomato, the SlMAP70-1 antibody recognize both endogenous SlMAP70 and over-expressed GFP-SlMAP70-1. The relative gray value corresponding to the endogenous SlMAP70 proteins (compared to actin control), which were lower in *SlMAP70-1* RNAi. **(b)** Western blot of protein extract from *N.benthamiana* leaves expressing of GFP-SlMAP70-1 (up), or non-transformed leaves (bottom). GFP antibody was used in lane 1. Three different SlMAP70-1 antibodies were used in lane 2-4. These SlMAP70-1 antibodies recognized both GFP-SlMAP70-1 and endogenous MAP70 of *N.benthamiana*. **(c)** Western blot of protein extract from *N.benthamiana* leaves expressing of CFP-SlMAP70-1 or SlMAP70-1 RNAi. The endogenous NtMAP70 level of *N.benthamiana* was reduced by the expression of SlMAP70 RNAi.

**Figure S7.**
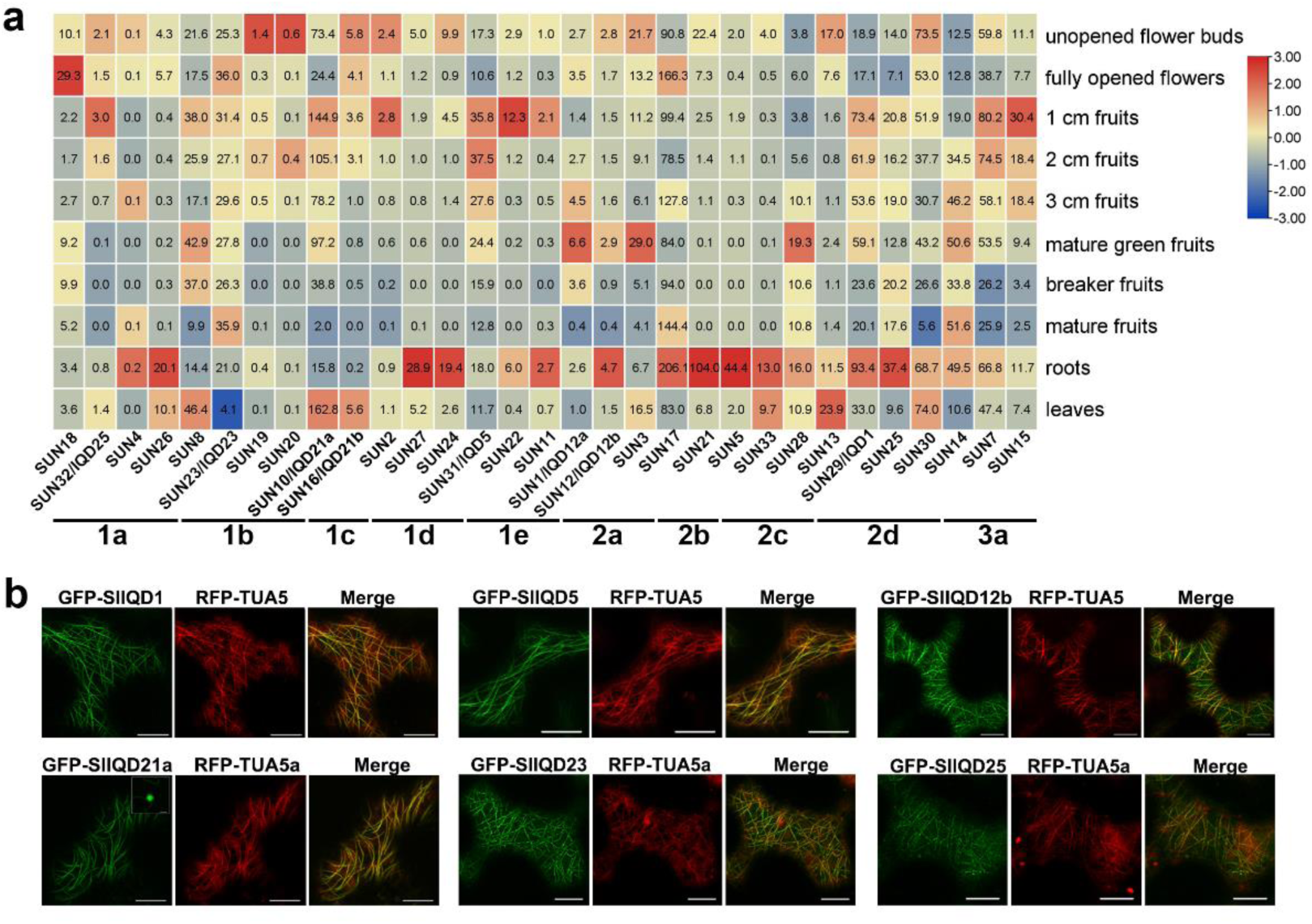
Subcellular localization and expression pattern of the SUN/SlIQD proteins. **(a)** Expression data of tomato *IQD* genes in different tissues and organs of tomato cultivar Heinz were obtained from the public RNA-seq database (http://ted.bti.cornell.edu/). It shows *SlIQD21a* gene highly expressed at fruit early stage (1 cm fruits). Bars and letters below the images indicate the phylogenetic clades of tomato IQD family (Huang *et al*., 2013). **(b)** Transient co-expression of *Pro35S:GFP-SlIQDs* with *Pro35S:RFP-TUA5* in *N. benthamiana* leaf epidermal cells. All SlIQD proteins selected from different phylogenetic clade were microtubule localized, except for SlIQD21a also labels nucleus. Bars: 10 μm.

**Figure S8.**
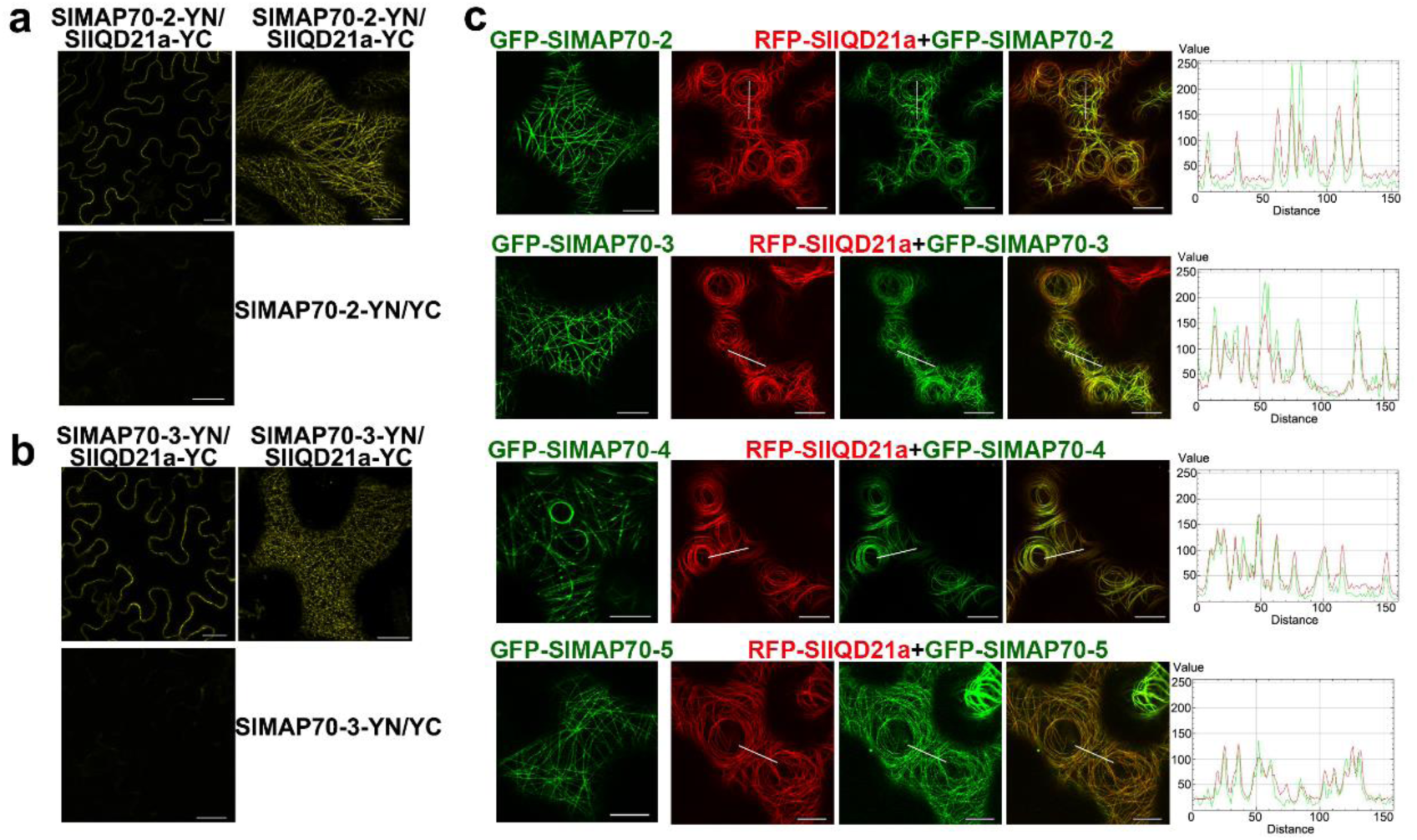
SlIQD21a is likely to interact with different SlMAP70 proteins in vivo. **(a-b)** BiFC assay showed that SlMAP70-2 and SlMAP70-3 proteins interact with SlIQD21a at microtubules in *N. benthamiana* epidermis cells. Single optical sections of YFP signals at equatorial plane (left) and spheric zone closed to plasma membrane (right) are shown. Free YN was used as a negative control. Bars: 10 μm. **(c)** Co-expression assays indicate tomato MAP70 family proteins co-localized with RFP-SlIQD21a fusion protein at cortical microtubules in *N. benthamiana* epidermis cells. Bars: 10 μm

**Figure S9.**
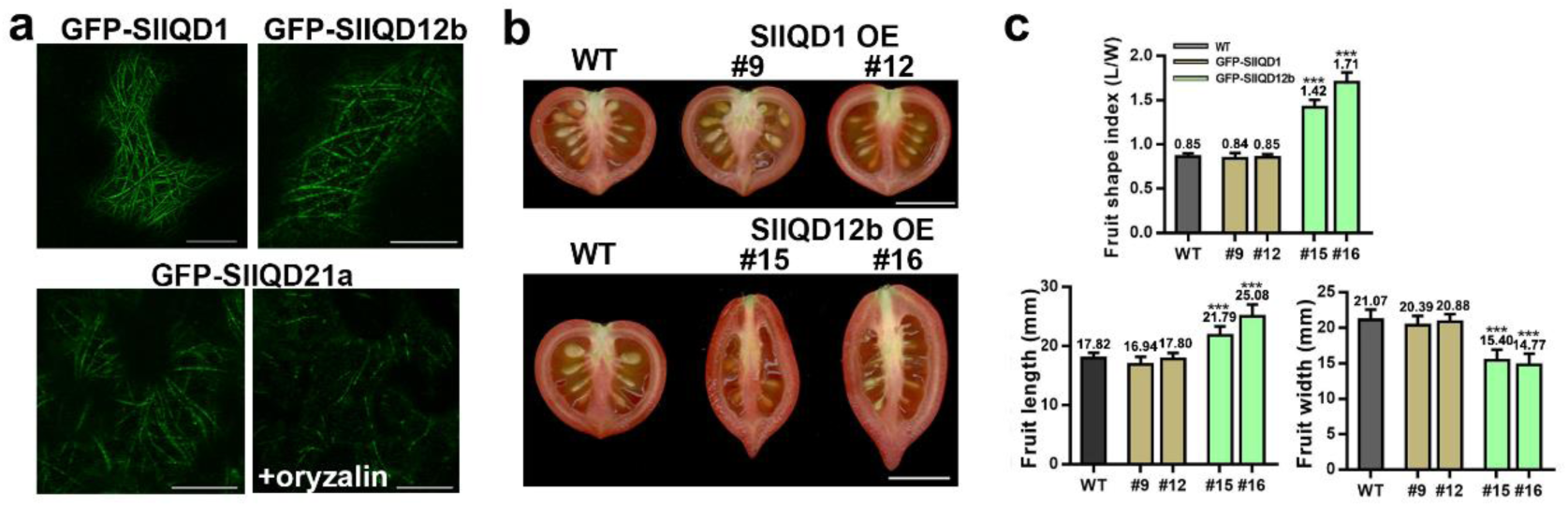
Fruit shape study of different SlIQD transgenic lines. **(a)** Subcellular localization of GFP-SlIQD1, GFP-SlIQD12b and GFP-SlIQD21a fusion proteins in tomato leave PCs under the control of the CaMV 35S promoter. All of them localized at microtubules, which can be depolymerized by the treatment with 100 nM oryzalin for 30min. Bars: 10 μm. **(b)** Phonotype of mature fruits of GFP-SlIQD1 and GFP-SlIQD12b overexpression plants. Bars: 1 cm. **(c)** Statistical analysis of the fruit length, width and shape index of IQD transgenic plants and WT. Data are shown as mean ± SD, n=30. ***, P < 0.001; *, P < 0.05 by Tukey’s test.

**Figure S10.**
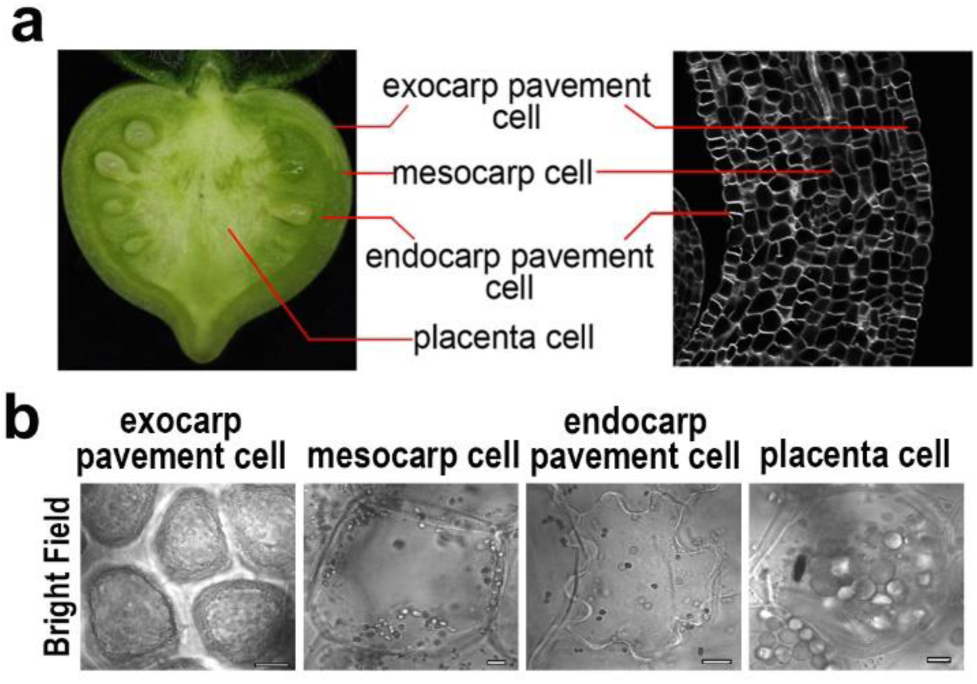
Schematic diagram of tomato fruit tissue. **(a)** A demonstration of different fruit tissues that were studied in (a). **(b)** Images of fruit cells from the tissues showed in (a). Bars: 10 μm.

**Figure S11.**
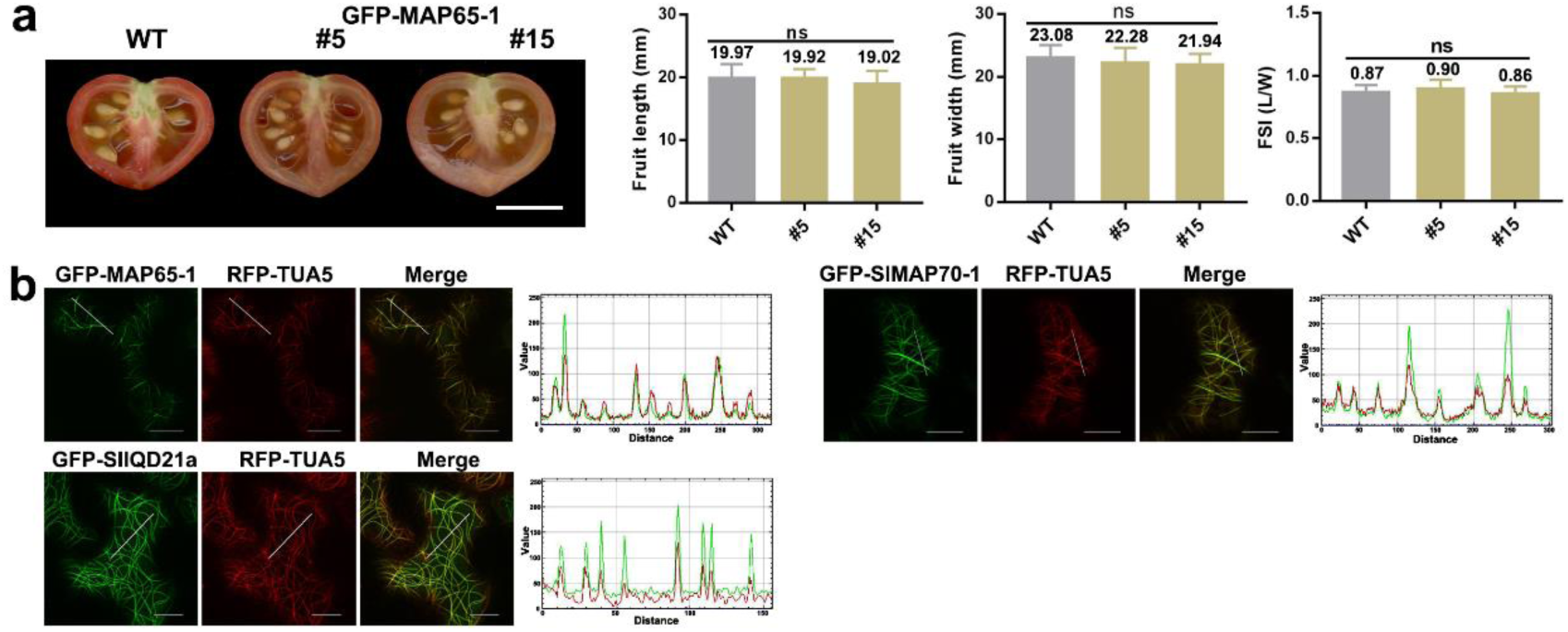
Effects of GFP-MAP65-1 on fruit shape and sub-cellular location analysis of MAP65-1, SlMAP70-1 and SlIQD21a proteins. **(a)** The expression of GFP-MAP65-1 fusion protein in ‘MicroTom’ tomato did not affect fruit morphology. Statistical analysis was performed by Tukey’s test with α < 0.05, n > 20. ns, no significance. Bar: 1cm. **(b)** Co-expression of GFP-MAP65-1, GFP-SlMAP70-1 and GFP-SlIQD21a proteins with RFP-TUA5 in *N. benthamiana* leaf epidermal cells, all of them co-localized with RFP-TUA5 labeled microtubules. Bars: 10 μm.

## Supplementary Movies

**Movie S1.** *N. benthamiana* leaf pavement cells expressing microtubule marker, GFP-TUB6.

**Movie S2.** *N. benthamiana* leaf pavement cells expressing GFP-SlMAP70-1 and RFP-SlIQD21a.

## Notes

### Competing Interest Statement

The authors have declared no competing interest.

