## Supplemental table S1 for "The microtubule-associated protein SlMAP70 interacts with SlIQD21 and regulates fruit shape formation in tomato"

**Supplementary Tables:**

**Table S1. Primers used for generating constructs**

| **Genes** | **Forward primers** | **Reverse primers** | **Plasmid** | **Purposes** |
| --- | --- | --- | --- | --- |
| SlMAP70-1 | cgcgtcgacATGGCGGAGGATGGAAGTG | cggaattctgTTATTGTGTGCGTGTTAACC | pMDC43 | Pro35S:GFP-SlMAP70-1 |
|  |  |  | pK7WGC2 | Pro35S:CFP-SlMAP70-1 |
|  |  |  | pGBKT7 | Y2H |
| SlMAP70-1 | gcgtcgacGAAGGGTCCATGCTGCTC | cggaattcAGCTGTTACCTTGGCACG | pHellsgate8 | SlMAP70-1 RNAi |
| SlMAP70-1/2 | gaatctaacagtgtagtttgAAACAGATAGAAATATGCCGTTTTAGAGCTAGAAATAGC | gctatttctagctctaaaacGGACAGCATTAGCAGCTGCCAAACTACACTGTTAGATTC | pTX | crispr |
| SlMAP70-1 | gaaccaattcagtcgacATGGCGGAGGATGGAAGTGTG | aagctgggtctagaccTTGTGTGCGTGTTAACCCACTT | YFPN | BiFC |
| SlMAP70-2 | aaaaagcaggctccATGTCGGAGTTTTCCGGCGAGT | agaaagctgggtgTTATTGAATGTTGCGTGTTAAC | pMDC43 | Pro35S:GFP-SlMAP70-2 |
|  |  |  | pGBKT7 | Y2H |
| SlMAP70-2 | gaaccaattcagtcgacATGTCGGAGTTTTCCGGCGAGT | aagctgggtctagaccTTGAATGTTGCGTGTTAACCCA | YFPN | BiFC |
| SlMAP70-3 | aaaaagcaggctccATGGCGGAGGTTTCCGGCGA | agaaagctgggtgTTATTGTGTGCCTCGCATTAATC | pMDC43 | Pro35S:GFP-SlMAP70-3 |
|  |  |  | pGBKT7 | Y2H |
| SlMAP70-3 | gaaccaattcagtcgacATGGCGGAGGTTTCCGGCGAT | aagctgggtctagaccTTGTGTGCCTCGCATTAATCCA | YFPN | BiFC |
| SlMAP70-4 | aaaaagcaggctccATGTCAGGCTTTTATGAAGTTTG | agaaagctgggtgTCATGATCCTTGTTGCTTCACT | pMDC43 | Pro35S:GFP-SlMAP70-4 |
|  |  |  | pGBKT7 | Y2H |
| SlMAP70-4/5 | gaatctaacagtgtagtttgATTTGCAGAACCAACTCAAGTTTTAGAGCTAGAAATAGC | gctatttctagctctaaaacCCTATCGAGTCCATTATTGCAAACTACACTGTTAGATTC | pTX | crispr |
| SlMAP70-5 | cgcgtcgacATGGCAGAGTTTGATGAATTTAG | gctctagaccTCAATATGTTTTGACCCTGACGG | pMDC43 | Pro35S:GFP-SlMAP70-5 |
|  |  |  | pGBKT7 | Y2H |
| SlIQD21a | cgcgtcgacATGGGCAAGAAAGGAAGTGG | gctctagaccTCAACTAAAATTATATCTCC | pMDC43 | Pro35S:GFP-SlIQD21a |
|  |  |  | pK7WGR2 | Pro35S:RFP-SlIQD21a |
|  |  |  | pGADT7 | Y2H |
| SlIQD21a | gaaccaattcagtcgacATGGGCAAGAAAGGAAGTG | aagctgggtctagaccACTAAAATTATATCTCC | YFPN | BiFC |
| MAP65-1 | cgggatccacATGGCAGCAGTAGATGATCA | gctctagaccCTATGGGGTGCTAGGAATAG | pMDC43 | Pro35S:GFP-MAP65-1 |
| TUA5 | aaaaagcaggctccATGAGGGAAATTATTAG | agaaagctgggtgTCAATAGTCTTCACCTTC | pK7WGR2 | Pro35S:RFP-TUA5 |
| SlIQD12b | cgcgtcgacATGGGGAAGAAAGGAAGCTG | cggaattctgCTAGTTGACAACTCCATTCA | pMDC43 | Pro35S:GFP-SlIQD12b |
|  |  |  | pGADT7 | Y2H |
| SlIQD1 | cgcgtcgacATGGGGAAGAAAGGAAGCTG | cggaattctgCTAGTTGACAACTCCATTCA | pMDC43 | Pro35S:GFP-SlIQD1 |
|  |  |  | pGADT7 | Y2H |
| SlIQD5 | gaaccaattcagtcgacATGGGTGTCTCTGGCAAATGG | aagctgggtctagaccATCAGCAGCCTGTTGTGAAACAG | pMDC43 | Pro35S:GFP-SlIQD5 |
|  |  |  | pGADT7 | Y2H |
| SlIQD21b | gaaccaattcagtcgacATGGGGAAGAAAGGAAGTGGT | aagctgggtctagaccCTAATTATAATTATATCTGTGAG | pGADT7 | Y2H |
| SlIQD23 | cgcgtcgacATGGGCAAAGCATCCAAATG | gctctagaccTTAATATCTGTGTCGAAGAC | pMDC43 | 35S:GFP-SlIQD23 |
|  |  |  | pGADT7 | Y2H |
| SlIQD25 | cgcgtcgacATGGGCAAAGCAATTAAGTG | gctctagaccTTAAAAGTCCCTTTCACCTT | pMDC43 | 35S:GFP-SlIQD25 |
|  |  |  | pGADT7 | Y2H |
| SlMAP70-1 peptide | actggtggacagcaaatgggtcgcggatccGATGGCAGAAGTTCAAGC | tggtggtgctcgagtgcggccgcaagcttgTTTTTCATCTTGATTTGA | pET28a | 6xHis-MAP70-1pep |
